# Artesunate Exploits Glutathione-Dependent Redox Vulnerability in Metastatic Lung Adenocarcinoma

**DOI:** 10.64898/2026.10.06.755294

**Authors:** Patricia Munteanu, Sophia Neuhaus, Paula Feldt, Vahan Serobyan, Anna Baginska, Jose Pedro Friedmann Angeli, Jürgen Becker, Alexander Schramm, Barbara M. Grüner

## Abstract

Metastatic dissemination remains the principal cause of mortality in lung adenocarcinoma. To identify metabolic vulnerabilities associated with metastasis, we used cell lines with distinct metastatic abilities derived from the *Kras^G12D^;p53^KO^*mouse model. Building on evidence of mitochondrial dysfunction in metastasis-derived cells, we performed an *in vivo* screen of a focused metabolic drug library using a multiplexed DNA-barcoding platform. This approach identified artesunate as a selective inhibitor of metastatic seeding, which was validated *in vivo* to reduce metastatic burden while leaving primary tumor growth unaffected. Mechanistically, artesunate induced oxidative stress, characterized by reactive oxygen species and lipid peroxidation. Transcriptomic and mechanistic profiling revealed altered redox homeostasis in metastasis-derived cells. Inhibition of glutathione synthesis specifically sensitized these cells to artesunate, resulting in extensive cell death, whereas primary tumor-derived cells remained resistant. These findings identify glutathione-dependent redox homeostasis as a metabolic vulnerability in metastatic cells that can be specifically targeted using artesunate.

## INTRODUCTION

Non–small cell lung cancer (NSCLC) is among the most common cancers worldwide and a major cause of cancer-related death (Hendriks et al., 2024). Despite advances in surgical resection, chemotherapy, targeted therapy, and immunotherapy, the prognosis for patients with advanced NSCLC remains poor, with a persistently low five-year survival rate (Li et al., 2023). The majority of patients eventually die from metastatic disease and more than 60% of lung cancer patients are diagnosed at advanced stages, when tumors have already disseminated and curative surgical intervention is no longer feasible (Osmani et al., 2018).

Metastasis is a complex, energy-intensive, multistep process requiring tumor cells to detach from the primary tumor, survive shear forces and oxidative stress in circulation, and successfully colonize distant organs (Fares et al., 2020). Metastasizing cells encounter high metabolic stress during dissemination and colonization, necessitating flexible adaptation of energy production and redox homeostasis (Bergers & Fendt, 2021). Single-cell RNA sequencing reveals that metastatic lesions display metabolic programs distinct from matched primary tumors, characterized by enriched mitochondrial oxidative metabolism (Davis et al., 2020). Metastasizing tumor cells experience elevated oxidative stress compared to established primary tumors, and this stress limits distant colonization (Piskounova et al., 2015). Furthermore, metastatic NSCLC cells exhibit altered mitochondrial functionality with lower membrane potential, higher ROS levels and heightened sensitivity to mitochondrial disruption (Chuang et al., 2021). Therefore, glutathione-dependent antioxidant pathways are increasingly recognized as selective regulators of metastatic progression. Gstt1, a member of the glutathione *S*-transferase superfamily, functions as a metastasis-specific driver without affecting primary tumor growth (Ferrer et al., 2024), and mitochondrial glutathione accumulation via SLC25A39 is required for efficient metastatic outgrowth (Yeh et al., 2025).

Despite this growing evidence of metabolic adaptations in metastatic lesions, it remains unclear whether these metastasis-associated metabolic states can be systematically exploited using pharmacological approaches. To uncover therapeutically actionable metastatic vulnerabilities, we assembled a curated metabolic drug library and tested its effects in matched primary tumor–derived (T_nonMet_) and metastasis-derived (Met) lung adenocarcinoma cell lines by adapting a multiplexed *in vivo* DNA barcoding platform (Grüner et al., 2016). This approach enabled high-throughput and unbiased evaluation of compounds with potential for therapeutic repurposing and their effects on metastatic seeding. Using this strategy, we identified artesunate as a selective inhibitor of metastatic seeding, acting through induction of reactive oxygen species, lipid peroxidation, and disruption of glutathione homeostasis. In addition, inhibition of glutathione synthesis markedly increased the sensitivity of Met cells to artesunate, whereas T_nonMet_ cells were largely unaffected, indicating a pronounced metastasis-specific reliance on redox homeostasis.

## RESULTS

### Multiplexed screening identifies artesunate as a metastasis-selective compound

To directly compare cell-intrinsic metabolic vulnerabilities between lung adenocarcinoma cells with distinct metastatic ability (Winslow et al., 2011), we adapted a multiplexed small-molecule barcoding–based *in vivo* screening platform previously developed for pancreatic cancer (Dorsch et al., 2021; Grüner et al., 2016) to lung cancer models. The screen was performed using a pair of lung adenocarcinoma cell lines generated *ex vivo* from the genetically engineered *Kras^G12D^;p53^KO^*;*Rosa26^Tom^*mouse model: 579T1 (T_nonMet_), derived from a primary tumor, and 579DLN (Met), derived from a matched metastasis (Chuang et al., 2021; Winslow et al., 2011). To identify metabolic dependencies associated with metastatic seeding, we applied this multiplexed *in vivo* screening platform to interrogate a curated library of metabolic compounds (Figure 1A). By comparing barcode abundance before injection and in cells recovered from the lungs 48 hours after injection, we quantified the effect of each compound on metastatic seeding. Parallel measurements of *in vitro* cell growth enabled us to distinguish metastasis-selective effects from general cytotoxicity. Using this approach, which primarily interrogates early stages of the metastatic cascade, including survival in the circulation and initial lung seeding (Grüner et al., 2016), we identified several compounds that selectively reduced metastatic seeding of T_nonMet_ cells, Met cells, or both (Figure 1B).

**Figure 1.**
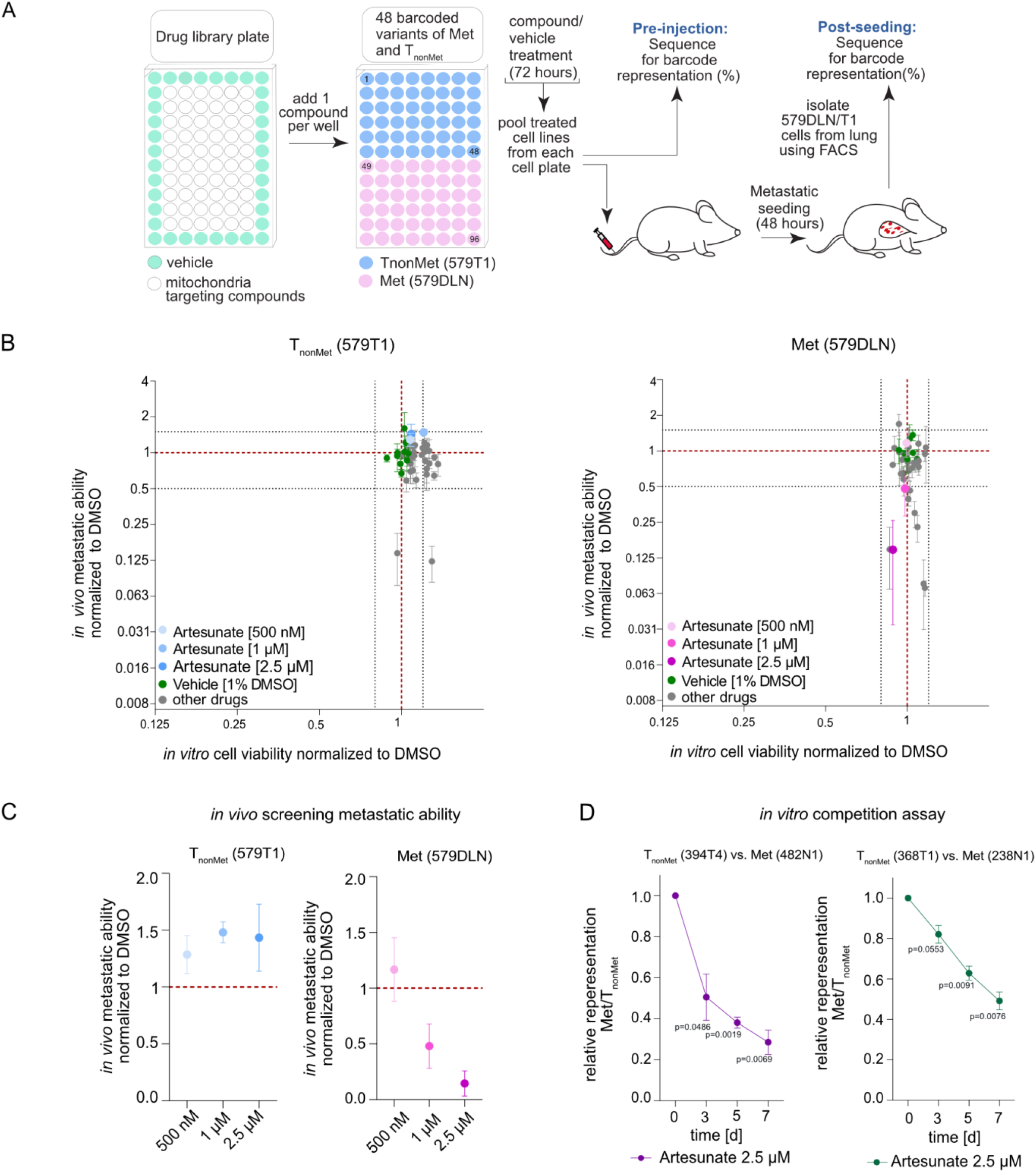
Multiplexed *in vivo* screening identifies artesunate as a selective inhibitor of metastatic seeding. **(A)** Schematic representation of the multiplexed barcoding-based screening workflow using a metabolic compound library. Barcoded T_nonMet_ (579T1) and Met (579DLN) cells were pretreated *in vitro* prior to intravenous injection, and metastatic seeding was quantified *in vivo* by barcode representation **(B)** Screening results for T_nonMet_ (579T1) and Met (579DLN) cell lines, plotting *in vitro* cell viability (confluence) on the x-axis against *in vivo* metastatic seeding normalized to control on the y-axis (mean ± SEM, *n* = 3). **(C)** Dose-dependent effects of artesunate on *in vivo* metastatic seeding of T_nonMet_ (579T1) and Met (579DLN) cells following 72 h *in vitro* pretreatment (mean ± SEM, *n* = 3). **(D)** *In vitro* competition assay of metastatic (238N1, 482N1) and non-metastatic (368T1, 394T4) cells treated with artesunate (2.5 µM). The ratio of metastatic to T_nonMet_ cells was normalized to the corresponding untreated control. Data are shown as mean ± SEM from *n* = 3 independent experiments. Statistical significance was assessed using Welch’s *t*-test comparing treated versus untreated conditions at each time point (Day 3, Day 5, and Day 7 relative to Day 0).

Among the tested compounds, the antimalarial compound artesunate emerged as one of the most prominent hits. Artesunate induced a strong, dose-dependent reduction in lung seeding of the Met (579DLN) cells, with minimal effects on *in vitro* growth and no detectable effect on lung seeding of the T_nonMet_ (579T1) cells (Figure 1C). This effect could not be explained by induction of apoptosis or reduced cell proliferation (Supplementary Figure 1A–B). To validate the selective sensitivity of Met cells to artesunate, we examined two additional T_nonMet_ (368T1, 394T4) and Met (238N1, 482N1) lung cancer cell lines derived from the same mouse model using *in vitro* competition assays. T_nonMet_ and Met cells were labeled with distinct fluorescent proteins (GFP or RFP) and jointly cultured in the presence of artesunate. Relative cell abundance was monitored over time by flow cytometry (Chuang et al., 2021). Across both additional models, artesunate treatment consistently reduced the proportion of Met cells compared to their T_nonMet_ counterparts (Figure 1D), though it did not induce significant changes in cell death or proliferation (Supplementary Figure 1A–B). These results confirm that artesunate preferentially impairs metastatic cells across multiple independent lung adenocarcinoma models.

### Artesunate selectively suppresses lung metastasis in a spontaneous metastasis model

Because the multiplexed screening approach primarily interrogates early metastatic seeding following intravenous injection and relies on *in vitro* pretreatment with artesunate, we next examined the effect of *in vivo* artesunate treatment in a spontaneous metastasis model that recapitulates the entire metastatic cascade. Following subcutaneous injection of Met (579DLN) cells, primary tumors were allowed to establish and spontaneously metastasize to the lungs. Upon tumor establishment, mice were treated daily with artesunate (100 mg/kg, i.p.) or vehicle for 16 days until control mice reached endpoint criteria (Figure 2A). Importantly, no metastatic lesions were detectable in the lungs at the time of treatment initiation, as confirmed by H&E analysis of a subset of mice sacrificed at this time point (Supplementary Figure 2A).

**Figure 2.**
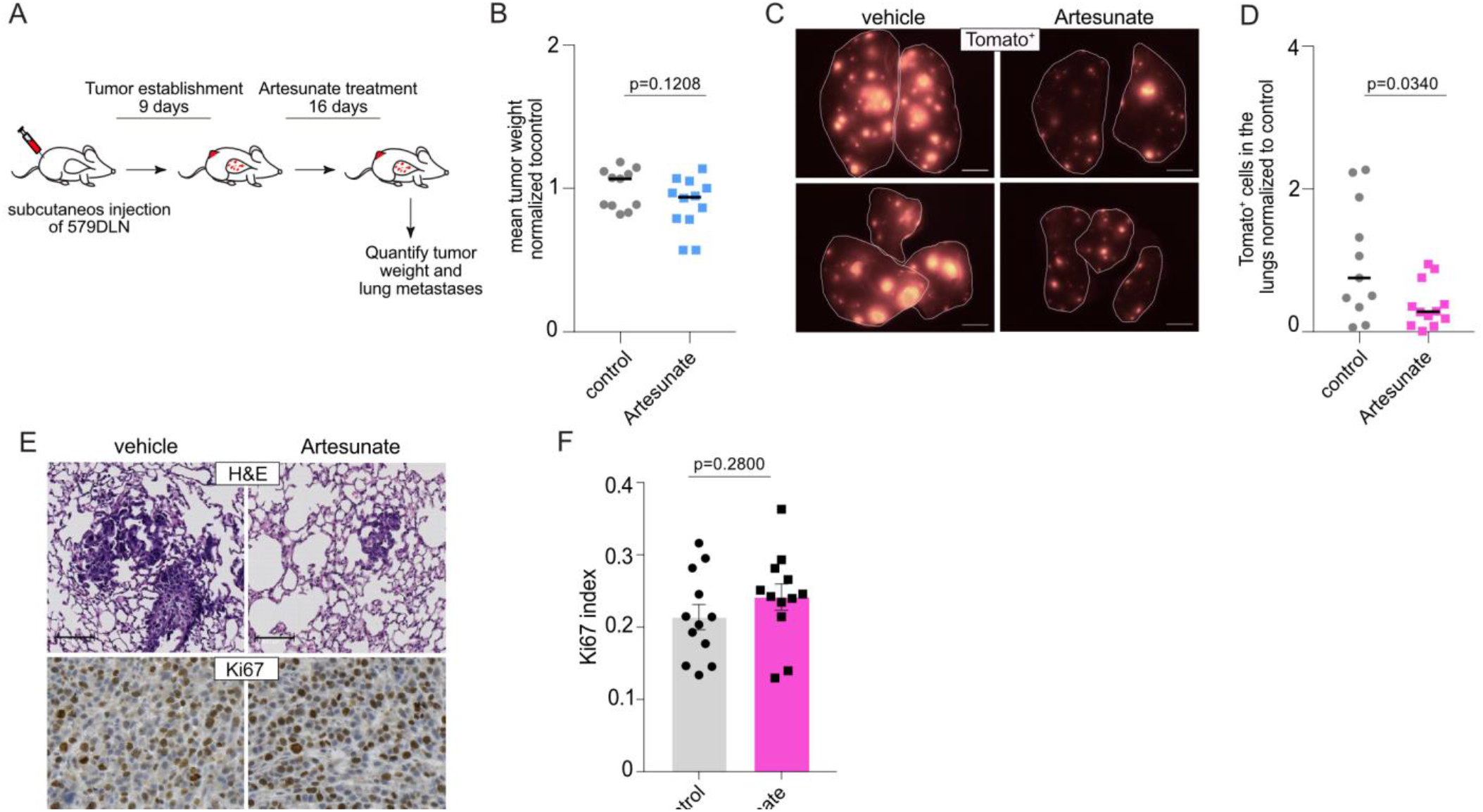
Artesunate selectively suppresses lung metastasis in a spontaneous metastasis model. **(A)** Schematic representation of the subcutaneous metastasis model and the corresponding experimental timeline. Mice were assigned to control (*n* = 11) or artesunate-treated (*n* = 12) groups. **(B)** Mean subcutaneous tumor weight per mouse in control and artesunate-treated groups (two tumors per mouse; control *n* = 11, artesunate *n* = 12). Statistical significance was assessed using Welch’s *t*-test following normality testing by the D’Agostino–Pearson test. **(C)** Representative fluorescence stereomicroscopy images of lungs from control and artesunate-treated mice showing metastatic burden. Scale bars, 1000 µm. **(D)** Quantification of Tomato⁺ cancer cells in the lungs by flow cytometry. Data were normalized to their respective control groups from two independent experiments. Each dot represents one mouse; the horizontal line indicates the mean. Statistical significance was assessed using Welch’s *t*-test following normality testing by the D’Agostino–Pearson test. **(E)** Representative H&E-stained lung sections and Ki67 immunohistochemistry (IHC) staining of tumors from vehicle- and artesunate-treated mice. Scale bars, 100 µm (H&E sections) and 50 µm (Ki67 staining). **(F)** Quantification of tumor cell proliferation based on Ki67 IHC. The Ki67 proliferation index was calculated as the ratio of Ki67⁺ nuclei to total tumor cell nuclei. For each mouse, two tumor sections were analyzed, with 10 randomly selected regions of identical size per section (20 regions of interest per mouse). Each data point represents the mean per mouse. Bars indicate mean ± SEM. Statistical significance was assessed using Welch’s *t*-test.

At the experimental endpoint, primary tumor weight and size were quantified, and metastatic burden was assessed by flow cytometric quantification of Tomato^positive^ tumor cells in the lungs. While primary tumor weight did not differ significantly between artesunate- and vehicle-treated groups (Figure 2B), artesunate treatment significantly reduced the number of tumor cells that metastasized to the lungs (p = 0.03) (Figure 2C–D). Quantification of Ki67 staining intensity revealed no significant difference in proliferation between artesunate- and vehicle-treated primary tumors (Figure 2E–F). These findings indicate that artesunate specifically reduces spontaneous lung metastasis independently of effects on primary tumor growth or proliferation. In addition, artesunate pretreatment did not significantly induce anoikis *in vitro* under the same conditions in multiple Met cell lines (Supplementary Figure 2B), suggesting that the reduced lung seeding is unlikely to result primarily from increased sensitivity to detachment-induced cell death (Dorsch et al., 2021).

### Transcriptional Profiling Reveals Differential Redox Responses and Glutathione Pathway Regulation upon Artesunate Treatment

To investigate the molecular basis of the metastasis-selective effects of artesunate, we performed RNA sequencing on Met and T_nonMet_ cells treated with artesunate or vehicle control for 48 hours. Pathway enrichment analysis of genes upregulated following artesunate treatment (Supplementary Figure 3A–B) revealed activation of the integrated stress response (ISR) and amino acid biosynthesis in both Met and T_nonMet_ cells, consistent with activation of a conserved transcriptional response to oxidative stress. However, distinct transcriptional adaptations were also evident. Met cells preferentially upregulated pathways involved in amino acid, folate, and cofactor metabolism, suggesting activation of metabolic pathways associated with adaptation to oxidative stress (Ducker & Rabinowitz, 2017; Yang & Vousden, 2016). In contrast, T_nonMet_ cells displayed increased enrichment of pathways related to endoplasmic reticulum function and lysosome biogenesis, consistent with activation of protein quality control and organelle homeostasis (Mancuso et al., 2021; N. D. Yang et al., 2014).

We next assessed pathways enriched among genes downregulated by artesunate (Figure 3A–B). Pathway enrichment analysis revealed a coordinated suppression of metabolic and signaling pathways in both cell lines. Shared downregulated pathways included glycolysis/gluconeogenesis, steroid biosynthesis, fatty acid metabolism, HIF-1 signaling, and TNF signaling, indicating a broad reduction in metabolic activity and stress-associated signaling. Beyond these common responses, Met cells preferentially exhibited decreased enrichment of pathways associated with cell-cycle progression, DNA replication, and extracellular matrix interactions, whereas T_nonMet_ displayed stronger suppression of PI3K-AKT and MAPK signaling as well as glutathione metabolism.

**Figure 3.**
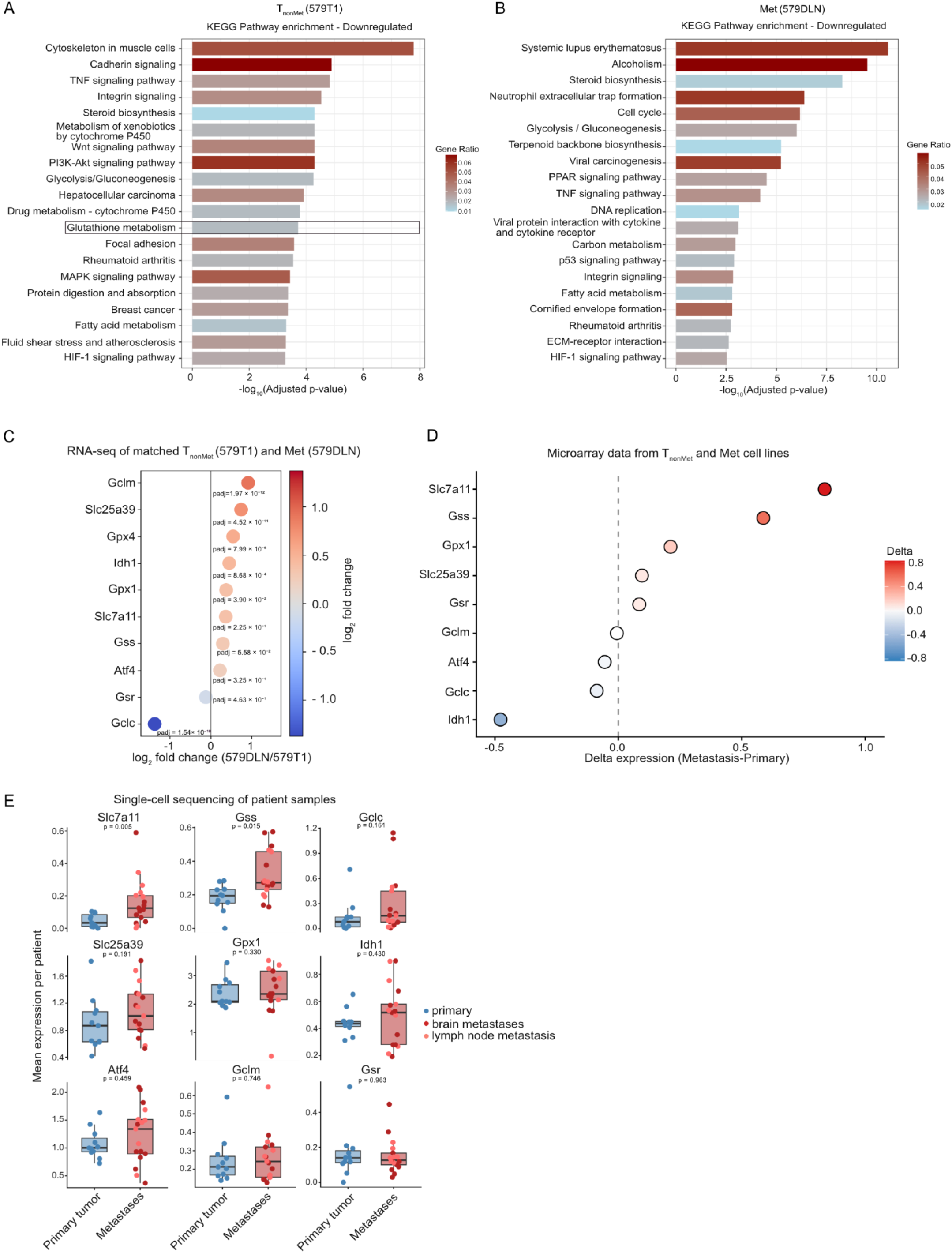
Transcriptional profiling reveals differential redox responses and glutathione pathway regulation upon artesunate treatment. **(A–B)** KEGG pathway enrichment analysis in 579T1 and 579DLN cells following 48 h treatment with 2.5 μM artesunate. RNA sequencing was performed using three independent biological replicates per condition. Bubble plots showing downregulated pathways based on bulk RNA sequencing. Downregulated pathways in 579T1 (A) and 579DLN (B) cells. The x-axis represents the −log_10_(adjusted p-value) of pathway enrichment, with larger values indicating greater statistical significance. Bar color indicates the gene ratio, representing the proportion of differentially expressed genes overlapping with each pathway. **(C)** Baseline expression of glutathione-related genes. Bubble plots show log₂ fold change of selected genes in Met (579DLN) relative to T_nonMet_ (579T1) cells under vehicle (0.5% DMSO) conditions, based on bulk RNA sequencing. **(D)** Expression of detected glutathione- and redox-related genes in murine lung adenocarcinoma cell lines derived from primary tumors (*n* = 14) or metastases (*n* = 9) (GEO dataset GSE26874; Winslow et al., 2011). Circles represent the difference in median expression (Δ = metastasis − primary) based on microarray data (Affymetrix Mouse Genome 430 2.0 Array). **(E)** Expression of glutathione-related genes in malignant epithelial cells from primary lung adenocarcinomas and metastatic lesions. Gene expression was analyzed using the published single-cell RNA sequencing dataset GSE131907. Malignant epithelial cells were identified based on the published cell type annotations. For each patient, the mean expression of the indicated genes was calculated, with each dot representing one patient. Primary tumors (blue, *n* = 11) were compared with metastatic lesions (red, *n* = 17), comprising brain metastases (dark red, *n* = 10) and lymph node metastases (light red, *n* = 7). Boxplots summarize the distribution of patient-level mean expression for each group. *P* values were calculated using a two-sided Wilcoxon rank-sum test on patient-level mean expression values.

Given the established role of ROS generation and oxidative stress in the biological activity of artesunate (Huang et al., 2022; Jiang et al., 2021), together with the activation of the integrated stress response observed in our transcriptomic analysis, we next focused on glutathione metabolism, a major regulator of cellular redox homeostasis and a key downstream effector of the integrated stress response during oxidative stress (Amiri et al., 2026). The differential enrichment of glutathione-related pathways between Met and T_nonMet_ cells suggested distinct redox-adaptive responses to artesunate and raised the possibility that these differences contribute to the distinct sensitivity of Met and T_nonMet_ cells to artesunate.

To determine whether the differential regulation of glutathione metabolism following artesunate treatment reflected differences in its basal activity, we examined the baseline expression of key genes involved in glutathione metabolism under untreated conditions. These included genes regulating cystine uptake (*Slc7a11*), glutathione synthesis (*Gclc, Gclm,* and *Gss*), glutathione recycling (*Gsr*), glutathione-dependent peroxide detoxification (*Gpx1*), mitochondrial glutathione import (*Slc25a39*), NADPH production (*Idh1*), and the integrated stress response (*Atf4*) (Amiri et al., 2026; Stockwell et al., 2017; Wang et al., 2021; C. Yang et al., 2014). Indeed, Met cells displayed elevated baseline expression of key genes involved in glutathione metabolism, suggesting a pre-existing adaptation to oxidative stress (Hayes & Dinkova-Kostova, 2014) (Figure 3C). These findings indicate that metastatic cells exist in a distinct redox state characterized by constitutively elevated expression of genes involved in glutathione metabolism.

To determine whether this phenotype extended beyond the 579DLN/579T1 pair, we analyzed microarray expression data of additional primary tumor- and metastasis-derived murine lung adenocarcinoma cell lines from the same *Kras^G12D^;p53^KO^* model (GSE26874). Metastasis-derived cell lines consistently exhibited enriched expression of glutathione metabolism genes, supporting the observation that metastatic cells are characterized by an altered basal redox state and enhanced glutathione-associated gene expression (Figure 3D).

To extend these findings to human disease, we analyzed publicly available single-cell RNA-sequencing data from primary lung adenocarcinomas and metastatic lesions (GSE131907). Consistent with our murine microarray results, SLC7A11 and GSS emerged as the most prominently expressed glutathione-associated genes in human tumors. Both genes were significantly upregulated in malignant epithelial cells from metastatic sites, including lymph node and brain metastases, compared with primary tumor cells, underscoring a conserved metabolic shift associated with metastatic progression (Figure 3E).

Although pathway enrichment analysis identified differential regulation of glutathione metabolism primarily in T_nonMet_ cells following artesunate treatment, analysis of individual differentially expressed genes revealed that several oxidative stress–responsive genes, including *Hmox1, Slc7a11, Cth,* and *Chac1* (Hayes & Dinkova-Kostova, 2014; Stockwell et al., 2017), were upregulated in both cell types. In addition, genes associated with mitochondrial function and metabolism, such as *Mthfd2, Aldh1l2, Cox6a,* and *Chchd10* (Ducker & Rabinowitz, 2017; Purandare et al., 2018; Spinelli & Haigis, 2018) were similarly induced (Supplementary Figure 3D–E). These findings align with prior reports that artesunate induces oxidative stress and mitochondrial dysfunction (Mercer et al., 2011; Zhang et al., 2018), indicating that both redox regulation and mitochondrial processes are central components of the artesunate response in NSCLC cells.

Together, these findings demonstrate that although both Met and T_nonMet_ cells activate a conserved oxidative stress response, including induction of the integrated stress response, they exhibit distinct regulation of glutathione metabolism. Importantly, the elevated basal expression of glutathione-associated genes in metastatic cells was consistently observed across murine models and human patient samples, suggesting a distinct redox state that may contribute to their differential response to artesunate. We therefore next investigated the functional role of ROS and glutathione-dependent redox homeostasis in mediating the anti-metastatic effects of artesunate.

### Artesunate perturbs cellular redox homeostasis by inducing oxidative stress and mitochondrial damage

To investigate how exactly artesunate suppresses metastasis, we next examined its effects on cellular redox homeostasis. Artesunate increased intracellular ROS levels in both Met and T_nonMet_ cell lines, and co-treatment with N-acetylcysteine (NAC), a ROS scavenger (Kalyanaraman, 2022), partially attenuated this artesunate-induced ROS accumulation, although the extent of rescue varied between cell lines (Figure 4A). To test whether ROS induction contributes to the reduced metastatic seeding observed following artesunate treatment, we used 579DLN cells and performed *in vivo* rescue experiments under matched pretreatment conditions. Cells were pretreated for 24 h with artesunate (2.5 µM), NAC (10 mM), or both, and injected intravenously into syngeneic 129/B6 F1 mice. Lung-seeded tumor cells were quantified 48 h later (Figure 4B). As expected, artesunate significantly reduced lung metastatic seeding, consistent with the results from our multiplexed screen (Figure 4C–D; one-way ANOVA, p = 0.03). NAC treatment alone had a modest effect, whereas NAC co-treatment only partially restored seeding in artesunate-treated cells, suggesting that oxidative stress may contribute to the artesunate-mediated inhibition of metastasis. Importantly, artesunate at 2.5 µM did not significantly affect cell viability under these pretreatment conditions as quantified by Annexin V and assessed by microscopy (Supplementary Figure 1A and 4A). Injected cells adhered and resumed growth when re-plated *in vitro* (Supplementary Figure 4B), confirming that reduced lung seeding did not result from compromised cell viability during pretreatment or the suspension phase prior to injection.

**Figure 4.**
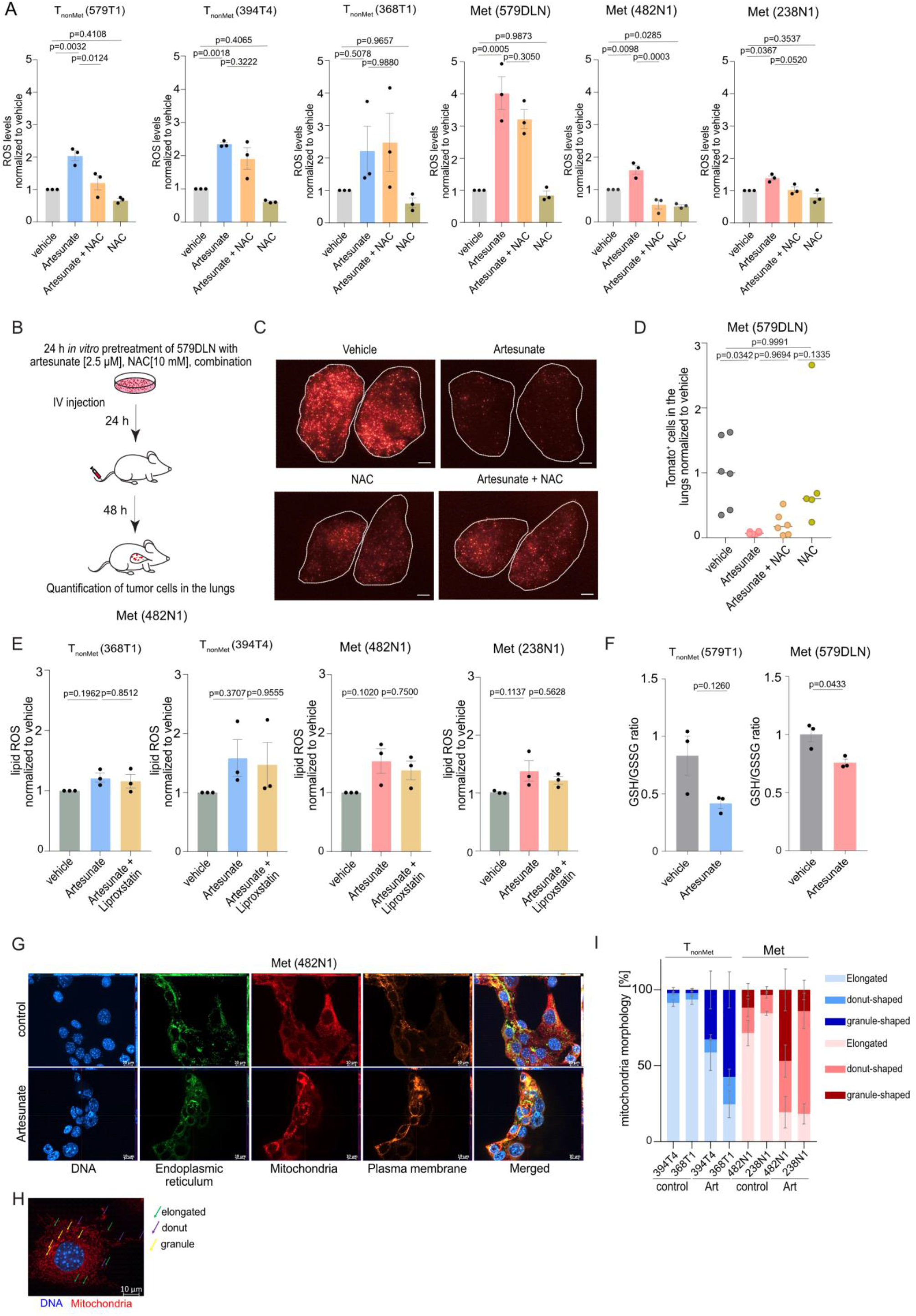
Artesunate perturbs cellular redox homeostasis by inducing oxidative stress and mitochondrial damage. **(A)** Quantification of ROS-high (FITC⁺) cells normalized to DMSO-treated controls following treatment with artesunate (2.5 μM), NAC (10 mM), or their combination for 24 hours. Data represent mean ± SEM from three independent experiments. Statistical significance was assessed by one-way ANOVA followed by Tukey’s multiple comparisons test. **(B)** Schematic overview of the *in vivo* lung seeding assay. Met (579DLN) cells were pretreated *in vitro* with artesunate (2.5 μM), NAC (10 mM), their combination, or vehicle for 24 h, followed by intravenous injection (1 × 10⁶ cells) into immunocompetent 129/B6 F1 mice. Lungs were harvested 48 h post-injection and analyzed by fluorescence stereomicroscopy and flow cytometry. **(C)** Representative fluorescence stereomicroscopy images of excised lungs 48 h post-injection showing Tomato⁺ tumor cells under the indicated treatments. Two lung lobes per condition are shown. Scale bars, 1000 µm. **(D)** Flow cytometric quantification of Tomato⁺ tumor cells in lung tissue 48 h post-injection. Each dot represents one mouse from two independent experiments. Data were normalized to the mean of the vehicle control for each experiment, and the normalized data from both experiments were combined. Statistical significance was assessed by one-way ANOVA followed by Tukey’s multiple comparisons test. **(E)** Lipid peroxidation 72 h after treatment with vehicle (0.5% DMSO), artesunate (2.5 µM), or artesunate plus Liproxstatin-1 (10 µM), measured using BODIPY 581/591 C11. The ratio of FITC to PE mean fluorescence intensity (MFI) was calculated and normalized to the respective vehicle control. Statistical significance was assessed by one-way ANOVA followed by Tukey’s multiple comparisons test. **(F)** Intracellular GSH/GSSG ratio was measured 24 h after artesunate treatment. Data are presented as the mean ± SEM of one representative experiment performed in triplicate. Three independent experiments were performed. Statistical significance was determined using Welch’s *t*-test. **(G)** Representative cell painting images of the Met 482N1 cell line under vehicle and artesunate-treated conditions (2.5 µM, 24 h). Individual channels show nuclei (blue), endoplasmic reticulum (green), mitochondria (red), and plasma membrane (orange), along with the corresponding merged image. Scale bars, 10 µm. **(H)** Representative images of nuclei and mitochondria, illustrating elongated, donut-shaped, and granular mitochondrial morphologies. Scale bars, 10 µm. **(I)** Quantification of mitochondrial morphology in T_nonMet_ and Met cell lines under vehicle and artesunate-treated conditions (2.5 µM, 24 h). Mitochondria were classified as elongated, donut-shaped, or granular, and are shown as percentage distribution per condition in the stacked bar graphs. Representative quantification from one of two independent experiments. In each experiment, five individual cells per condition were analyzed. Data represent mean ± SEM.

Consistent with the increase in intracellular ROS, artesunate also induced lipid peroxidation, although the magnitude of this effect varied across the tested cell lines (Figure 4E). T_nonMet_ cells showed greater heterogeneity, with little to no increase in 368T1 cells but a more pronounced response in 394T4 cells, whereas both Met cell lines exhibited a consistent increase in lipid peroxidation. Co-treatment with Liproxstatin-1, a radical-trapping antioxidant that inhibits lipid peroxidation (Zilka et al., 2017), partially attenuated this phenotype (Figure 4E). Because glutathione plays a central role in cellular ROS detoxification and is required for GPX4-dependent reduction of lipid peroxides (Wahida & Conrad, 2025), we next assessed glutathione redox homeostasis. Artesunate altered intracellular GSH and GSSG levels in both Met and T_nonMet_ cells, resulting in a reduced GSH/GSSG ratio, indicative of impaired glutathione-dependent antioxidant capacity (Figure 4F). Despite the disruption of glutathione redox homeostasis, GPX4 protein abundance remained unchanged following artesunate treatment in both Met and T_nonMet_ cells (Supplementary Figure 4C), suggesting that the observed lipid peroxidation is unlikely to result from reduced GPX4 expression.

Using cell painting-based analysis, we next assessed mitochondrial structure in Met and T_nonMet_ cells following artesunate treatment (Figure 4G–H). Without intervention, mitochondria in all cell lines predominantly displayed an elongated morphology, consistent with an intact network. Upon artesunate treatment, mitochondria exhibited a shift toward donut-shaped and granular morphologies, reflecting intermediate and fragmented states, respectively (Figure 4H–I). This shift was consistently observed in both Met cell lines that could be tested (those without endogenous tdTomato expression), which showed a pronounced reduction in elongated mitochondria and an increase in granular structures. In contrast, T_nonMet_ cell lines displayed greater variability, with 368T1 showing a more evident redistribution toward granular structures, whereas 394T4 retained a higher proportion of elongated mitochondria under the same conditions (Figure 4I). Together, these data indicate that artesunate treatment is associated with changes in mitochondrial structure, with a more pronounced effect observed in Met cells.

Overall, these findings demonstrate that artesunate induces oxidative stress in lung cancer cells and affects mitochondrial function. However, the comparable induction of ROS in both cell types suggested that intrinsic differences in redox homeostasis, rather than ROS levels alone, may underlie the increased sensitivity of metastatic cells.

### Metastasis-derived lung cancer cells exhibit elevated basal oxidative stress and altered glutathione homeostasis

Next, we asked whether the elevated expression of glutathione-associated genes identified in Met cells by transcriptomic and microarray analyses (Figure 3E–F) is accompanied by alterations in redox homeostasis that could explain their increased sensitivity to artesunate. Functionally, baseline measurements indicated a general trend toward higher intracellular ROS levels in metastatic cell lines relative to primary tumor-derived cell lines, except for 579DLN, which displayed lower baseline ROS (Figure 5A). In contrast, total glutathione (GSH) levels were generally lower in metastatic than in primary tumor-derived cell lines, although this reduction was more variable across models (Figure 5B). These findings suggest that Met cells exist in a more oxidized basal state and depend more heavily on glutathione-mediated redox buffering, thereby increasing their vulnerability to further ROS induction.

**Figure 5.**
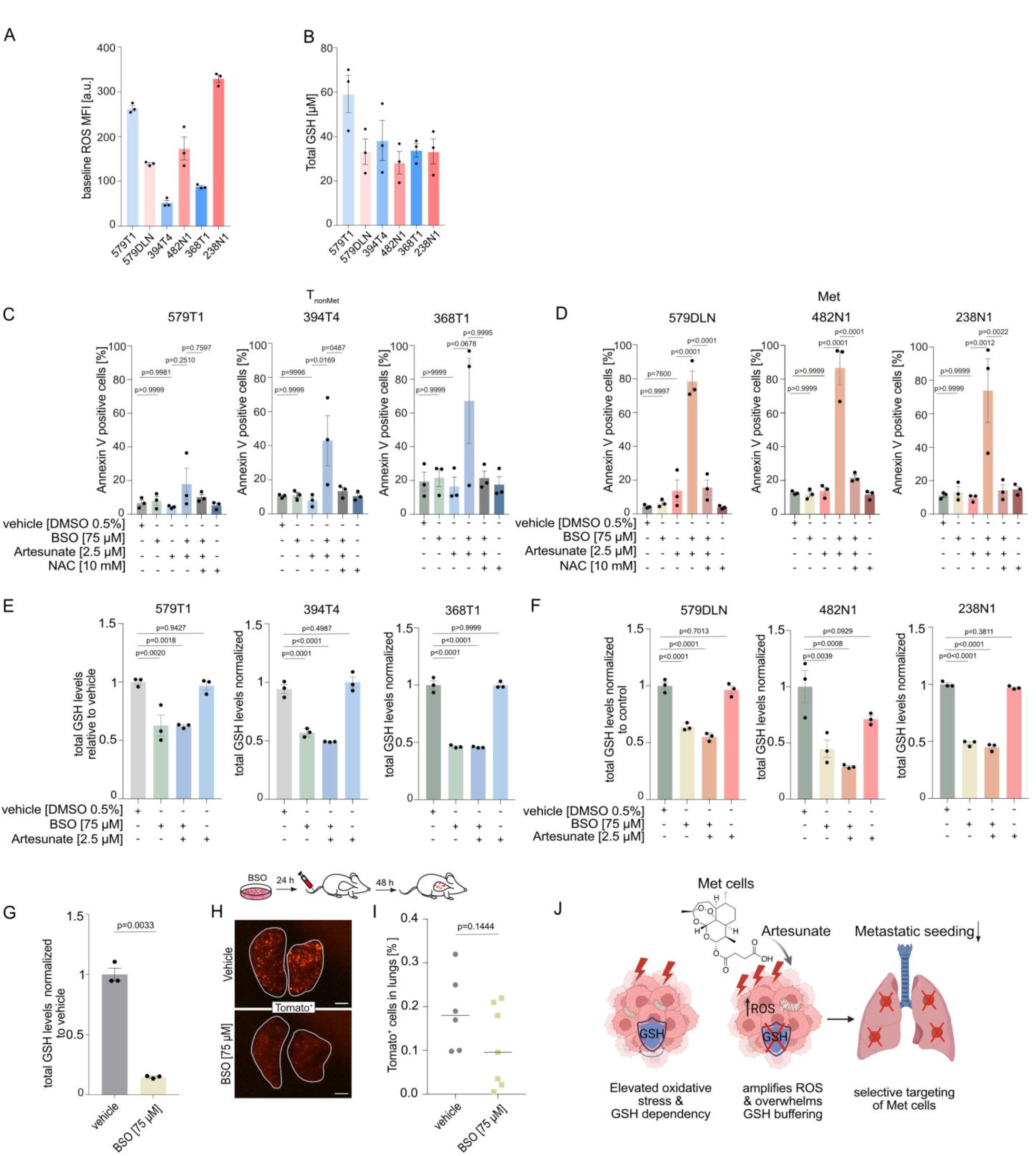
Metastasis-derived lung cancer cells exhibit elevated basal oxidative stress and altered glutathione homeostasis. **(A)** Baseline ROS MFI levels in Met and T_nonMet_ cells from a representative experiment. Data are shown as mean ± SEM of technical replicates. Similar results were obtained in all three independent experiments. **(B)** Baseline GSH levels in Met and T_nonMet_ cell lines. Data represent mean ± SEM from *n* = 3 independent biological replicates. **(C-D)** Flow cytometric quantification of Annexin V–positive cells in T_nonMet_ (C) and Met (D) cells 72 h following treatment with DMSO, artesunate (2.5 μM), artesunate + BSO (75 μM), or artesunate + BSO + NAC (10 mM). Early and late apoptotic populations were combined as total Annexin V–positive cells. Data represent mean ± SEM from three independent experiments. Statistical significance was assessed by one-way ANOVA followed by Tukey’s multiple comparisons test. **(E-F)** Total glutathione (GSH) levels in T_nonMet_ (E) and Met (F) cells 6 h after treatment with DMSO, artesunate (2.5 µM), BSO (75 µM), or their combination. Each dot represents one technical replicate (mean ± SEM). Statistical significance was assessed by one-way ANOVA followed by Dunnett’s multiple comparisons test. One representative experiment of three independent biological replicates is shown. **(G)** Total glutathione (GSH) levels in 579DLN cells 24 h after treatment with BSO (75 µM). Data are presented as mean ± SEM. One representative experiment of three independent biological replicates is shown. Statistical significance was assessed using a two-tailed Welch’s *t*-test. **(H)** Representative fluorescence stereomicroscopy images of lungs from mice injected with control- or BSO-pretreated 579DLN Tomato⁺ cells, showing Tomato⁺ cells seeded in the lungs. Scale bars, 1000 µm. **(I)** Quantification of Tomato⁺ 579DLN cells in lung tissue by flow cytometry (FACS) 48 h after intravenous injection of control- or BSO-pretreated (75 µM) cells. Data were normalized to the control group. Each dot represents one mouse, and the black line indicates the median. Statistical significance was determined using Welch’s *t*-test. **(J)** Working model summarizing the mechanism of artesunate action in Met cells. Met cells rely on glutathione-mediated redox buffering to maintain survival and support metastatic seeding. Artesunate induces ROS, overwhelming the limited antioxidant capacity of these cells. Depletion of glutathione further sensitizes Met cells to artesunate and reduces metastatic lung seeding. Schematic created with BioRender.com.

To functionally test this hypothesis, we combined artesunate with the glutathione synthesis inhibitor buthionine sulfoximine (BSO) (Griffith & Meister, 1979) in Met and T_nonMet_ cells (Figure 5C–D). While artesunate or BSO alone had minimal effects on cell viability, the combination induced near-complete cell death in all Met cells (Figure 5D). In contrast, T_nonMet_ cells exhibited heterogeneous responses: 579T1 cells remained largely unaffected, 394T4 cells displayed only moderate sensitivity, and 368T1 cells showed a non-significant trend toward increased sensitivity (Figure 5C). Importantly, co-treatment with N-acetylcysteine (NAC) fully rescued artesunate + BSO-induced cell death, confirming that cell death was mediated by oxidative stress (Figure 5C–D). Liproxstatin-1 similarly restored viability, indicating that lipid peroxidation contributes to this redox-dependent cell death (Supplementary Figure 5A–B).

To confirm effective inhibition of glutathione synthesis, intracellular glutathione levels were measured at an early time point before the onset of detectable cell death in the combination treatment. At this early time point, BSO significantly reduced intracellular glutathione levels in both Met and T_nonMet_ cells, and this effect was slightly enhanced by co-treatment with artesunate, whereas artesunate alone had no significant effect (Figure 5E–F). Notably, despite comparable glutathione depletion in both cell types, only Met cells underwent pronounced cell death upon combination treatment, confirming a greater dependence on glutathione synthesis under oxidative stress.

Given the increased dependence of Met cells on glutathione-mediated redox buffering, we next asked whether this requirement also contributes to metastatic seeding. Pretreatment of 579DLN cells with BSO (75 μM, 24 h) significantly reduced intracellular glutathione levels (Figure 5G) without affecting cell growth *in vitro* (Supplementary Figure 5C). Importantly, upon intravenous injection, glutathione-depleted cells showed a trend toward reduced lung seeding compared with controls (p = 0.14, Figure 5H–I). Thus, glutathione appears to contribute to efficient metastatic seeding in Met cells.

Together, these findings suggest that Met cells are more reliant on glutathione-mediated redox buffering when challenged with oxidative stress. Artesunate exploits this vulnerability by inducing oxidative stress in cells with limited redox capacity, thereby promoting redox imbalance when glutathione levels are further reduced. Glutathione depletion showed a trend toward reduced metastatic lung seeding, suggesting that maintenance of glutathione pools may support metastatic seeding and could contribute to the anti-metastatic effects of artesunate (Figure 5J). Further experiments using conditions that achieve more extensive glutathione depletion will be required to determine whether more extensive glutathione depletion can further reduce metastatic seeding.

## DISCUSSION

Metastatic dissemination is the primary driver of cancer-related mortality (Lambert et al., 2017). Clinically, this challenge is reflected in the poor prognosis of metastatic non–small cell lung cancer (NSCLC), which carries a 5-year survival rate of approximately 4%, underscoring the urgent need to identify metastasis-specific vulnerabilities (Belaroussi et al., 2023). Despite its clinical impact, the successful establishment of metastasis is a highly inefficient process, requiring disseminated tumor cells to overcome multiple barriers (Lambert et al., 2017). Increasing evidence indicates that metastatic seeding depends not only on genetic and transcriptomic alterations but also on adaptive metabolic reprogramming that enables survival in foreign microenvironments with distinct nutrient availability and oxidative pressures (Bergers & Fendt, 2021). For example, melanoma cells rewire antioxidant metabolism to endure oxidative stress during systemic dissemination (Piskounova et al., 2015), and lung cancer brain metastases exhibit enhanced glutathione metabolism and antioxidant defenses, reflecting adaptation to elevated redox stress (Liu et al., 2021).

In this study, we utilized cell lines derived from non-metastatic primary tumors (T_nonMet_) and metastases (Met) from the *Kras*^G12D^;*Trp53*^KO^ model that retain their intrinsic metastatic properties (Winslow et al., 2011). We previously demonstrated that these cells possess distinct mitochondrial functionality, with Met cells having lower mitochondrial membrane potential and higher ROS levels, which renders them specifically vulnerable to anti-mitochondrial targeting (Chuang et al., 2021). However, anti-mitochondrial targeting to prevent metastasis so far is not achievable safely with established compounds, particularly not in long-term treatment settings. Therefore, we interrogated lung adenocarcinoma cells for metastasis-specific vulnerabilities by adapting a previously established multiplexed *in vivo* screening platform (Dorsch et al., 2021; Grüner et al., 2016). With this, we screened a focused library of compounds targeting diverse aspects of mitochondrial metabolism for their effects on T_nonMet_ and Met cells. In this setting, artesunate emerged as the most consistent hit, selectively impairing metastatic seeding of Met cells across all tested *in vivo* models.

Artesunate, an FDA-approved drug for the treatment of severe malaria (Thomas et al., 2023), has previously been reported to suppress tumor growth and metastatic dissemination in several cancer types, including Merkel cell carcinoma, colorectal and renal cell carcinoma (Jeong et al., 2015; Li et al., 2007; Sarma et al., 2020). In addition, artesunate inhibited metastatic traits in lung cancer models *in vitro* and in chorioallantoic membrane assays, an effect that was largely attributed to modulation of matrix metalloproteinases (Rasheed et al., 2010). Our findings extend these observations by demonstrating selective activity against Met cells *in vivo* in both immunocompromised and immunocompetent settings and by providing evidence for an association with glutathione-dependent redox homeostasis.

As previously demonstrated, Met cells exhibited elevated baseline levels of reactive oxygen species compared with T_nonMet_ cells, suggesting that they operate closer to the threshold of tolerable oxidative stress (Chuang et al., 2021). In line with this vulnerability, we find that Met cells show increased expression of multiple components of the glutathione antioxidant axis, including *Slc7a11, Gss, Gpx1, Gsr*, and *Slc25a39*, indicating an enhanced reliance on glutathione-dependent redox defence mechanisms. This transcriptional program was robust across all Met cell lines analyzed and was further supported by publicly available single-cell RNA-sequencing data from patients with lung adenocarcinoma, demonstrating that this redox-associated phenotype is conserved in human metastatic disease. Together, these findings strengthen the generalizability of our observations and highlight glutathione-mediated antioxidant buffering as a recurrent feature of metastatic cells.

Consistent with this glutathione-associated phenotype, co-treatment with artesunate and the glutathione synthesis inhibitor BSO resulted in near-complete cell death in Met cells, whereas T_nonMet_ cells were largely resistant or only partially affected. Rescue by N-acetylcysteine (NAC) and Liproxstatin-1 further demonstrated that this lethality is driven by oxidative stress and lipid peroxidation (Fan et al., 2021; Pedre et al., 2021). Importantly, although BSO reduced intracellular glutathione levels in both Met and T_nonMet_ cells, pronounced cell death occurred selectively in Met cells, indicating a greater reliance on glutathione-mediated redox buffering. Together, these findings support a model in which Met cells exhibit heightened sensitivity to oxidative stress, enabling artesunate to selectively exploit a glutathione-dependent vulnerability at concentrations not yet affecting non-metastatic cells. Notably, glutathione depletion showed no effect on *in vitro* proliferation but a trend toward reduced metastatic lung seeding of Met cells, suggesting that glutathione dependence may extend beyond protection from pharmacologically induced oxidative stress. Further studies using more extensive glutathione depletion will be required to determine whether the trend toward reduced metastatic seeding is enhanced under conditions of more pronounced glutathione depletion.

Consistent with this model, oxidative stress buffering and glutathione-dependent metabolic adaptations have been implicated in metastatic progression across multiple cancer types, including melanoma, colorectal cancer, and breast cancer (Biondini et al., 2024; Laoukili et al., 2022; Piskounova et al., 2015). Furthermore, loss of the mitochondrial glutathione transporter *Slc25a39* has been shown to impair metastatic colonization without affecting primary tumor growth, highlighting a critical role for mitochondrial glutathione in metastatic progression (Yeh et al., 2025). Together, these findings suggest that glutathione-dependent redox homeostasis constitutes an actionable vulnerability of metastatic cells.

Importantly, artesunate is an FDA-approved antimalarial agent that has been evaluated in early-phase clinical studies across multiple malignancies, including colorectal cancer, advanced solid tumors, and metastatic breast cancer, where it demonstrated favorable tolerability together with preliminary evidence of anti-tumor activity (Deeken et al., 2018; Konig et al., 2016; Krishna et al., 2015; von Hagens et al., 2019). These clinical studies, together with our findings, provide a strong rationale for further evaluating FDA-approved artesunate as a therapeutic strategy to target redox vulnerabilities in metastatic disease. In conclusion, our study not only identifies oxidative stress regulation as a bottleneck in lung adenocarcinoma metastasis but also provides artesunate as a promising repurposing candidate that exploits glutathione homeostasis in metastasis-derived lung cancer cells. By targeting this adaptive redox program, artesunate reveals a therapeutically actionable metabolic vulnerability and provides a rationale for further translational investigation.

## METHODS

### Cell lines

Mouse NSCLC cell lines were derived from primary tumors (T_nonMet_: 368T1, 394T4, 579T1) and metastatic lesions (Met: 238N1, 482N1, 579DLN) arising in genetically engineered mice harbouring the *Kras*^LSL-G12D/+^;*Trp53*^flox/flox^ (*Kras*^G12D^; *Trp53*^KO^) or *Kras*^LSL-G12D/+^;*Trp53*^flox/flox^; Rosa26^LSL-Tomato/+^ (*Kras*^G12D^;*Trp53*^KO^;R26^Tomato^) alleles (Winslow et al., 2011). The human HEK293T cell line was obtained from ATCC (ATCC Cat. No. CRL-3216, RRID: CVCL_0063).

### Cell culture

All cell lines were cultured at 37 °C and 5% CO₂, in DMEM (Gibco), supplemented with 10% FBS (VWR/Biowest), 50 U/mL penicillin, and 50 μg/mL streptomycin (Thermo Fisher Scientific). Cell lines were maintained in culture for no more than 1 month, after which a new vial was thawed. Cells were frozen in DMEM medium containing 20% FBS and 10% DMSO (Carl Roth). Passaging was performed once the cells reached 70–80% confluency by trypsinization, and a fraction of detached cells was reseeded into fresh cell culture medium. All cell lines used in this study were regularly tested for mycoplasma contamination.

### Reagents

Compounds used in the barcoding screen are listed in Supplementary Table 1A. All compounds were obtained as a focused custom-picked library from MedChemExpress. For *in vitro* experiments, commercially available compounds were used, including artesunate (Biorbyt, orb1480488), L-buthionine-(S,R)-sulfoximine (BSO) (MedChemExpress, HY106376), N-acetyl-L-cysteine (NAC) (Sigma-Aldrich, A9165-25G), and Liproxstatin-1 (Hölzel-Biotech, MOLN-16796). All drugs were dissolved in DMSO (Carl Roth, A994.2) so that the final concentration of DMSO in culture medium did not exceed 1%, and vehicle control samples received DMSO alone at the same concentration. For the *in vivo* treatment experiments, artesunate (Hölzel-Biotech, MOLN-M16406) was used at 100 mg/kg dissolved in saline with 2.5 % NaHCO₃ (Sigma-Aldrich, S6014).

### *In vitro* experiments

Unless stated otherwise, all *in vitro* experiments were performed in at least three independent biological experiments, each with three technical replicates.

### Confluency measurements

Cell confluency was monitored using the NyOne automated live-cell imager, and image analysis was performed with YT-Software (Synentec). Cells were seeded in 96-well or 12-well plates at an initial confluency of approximately 20–40%. Confluency was measured before treatment (day 1) and at subsequent time points following treatment. For the multiplexed barcoding platform, confluency values were normalized to the initial (day 1) measurement and to untreated controls. For artesunate and BSO treatment experiments, data are presented as the percentage of confluency relative to the initial measurement over time.

### Annexin V assay

Cell death was assessed using Annexin V–FITC staining (BD Pharmingen™, 556420) according to the manufacturer’s instructions. Cells were pretreated with artesunate (2.5 µM) for 72 h, harvested, and stained with Annexin V–FITC. Apoptotic cells were quantified using the BD FACSCelesta™ Analyzer (Becton, Dickinson and Company). Data were analyzed with FlowJo® software, and the percentage of Annexin V–positive cells was plotted for the control and artesunate-treated samples.

### Competition assay

T_nonMet_ (368T1, 394T4) and Met (482N1, 238N1) cell lines were stably labeled with GFP or RFP, mixed at a 1:1 ratio, and seeded in 6-well plates as described before (Chuang et al., 2021). Cells were treated with artesunate (2.5 μM) for 7 days, with drug replenished every 2– 3 days. Relative population dynamics were assessed by flow cytometry at the indicated time points and analyzed using FlowJo® software.

### Glutathione depletion and rescue assay

Cells were seeded at 1 × 10⁵ cells per well in 12-well plates. Twenty-four hours after seeding, cells were treated for 72 h with vehicle control (0.5% DMSO), artesunate (2.5 μM), L-buthionine-(S,R)-sulfoximine (BSO; 75 μM), N-acetylcysteine (NAC; 10 mM), artesunate + BSO, or artesunate + BSO + NAC (rescue condition).

Following treatment, cells were harvested and stained with Annexin V–FITC for 15 min in the dark. DAPI (5 μg/mL) was used to identify dead cells. Samples were measured within 1 h on a flow cytometer (BD FACSCelesta™, BD Biosciences) using a 488-nm laser, and at least 10,000 singlet events were recorded per sample. Data were analyzed using FlowJo® software. Cell death was quantified as the percentage of Annexin V–positive cells, including both early and late apoptotic populations.

### ROS assay

Intracellular ROS levels were measured using the fluorescent probe CM-H₂DCFDA (Thermo Fisher Scientific). Cells were seeded at 1 × 10⁵ cells per well in 12-well plates and allowed to adhere overnight. Where indicated, cells were co-treated with artesunate (2.5 µM) and N-acetylcysteine (NAC; 10 mM) to assess ROS scavenging.

Following treatment, cells were harvested by trypsinization, centrifuged, and resuspended in pre-warmed PBS containing 1 μM CM-H₂DCFDA. Staining was performed for 30 min at 37 °C in the dark. Hydrogen peroxide (H₂O₂) was used as a positive control, and unstained cells served as negative controls.

After staining, cells were washed and resuspended in PBS containing 2% FBS and DAPI for viability discrimination. Samples were analyzed by flow cytometry (BD FACSCelesta™, BD Biosciences) using FITC detection. Doublets and dead cells were excluded based on FSC/SSC parameters and DAPI positivity. ROS levels were quantified as the percentage of FITC-positive viable cells and normalized to DMSO-treated controls. Data were analyzed using FlowJo® software.

### Lipid peroxidation assay

For measurement of lipid ROS, metastasis-derived Met (238N1, 482N1) and T_nonMet_ (368T1, 394T4) cell lines were seeded in 12-well plates and treated with artesunate (2.5 μM) or with artesunate and Liproxstatin-1 (10 µM) for 72 h.

Following treatment, C11-BODIPY 581/591 (Cayman Chemical, #CAY27086-1) was added at a final concentration of 5 μM and incubated for 30 min at 37 °C in a humidified incubator with 5% CO₂. Cells were then harvested, centrifuged at 300 × g for 3 min at 4 °C, washed with PBS, and resuspended in a 200 μL FACS buffer for flow cytometric analysis.

Data were analyzed using FlowJo® software. Lipid ROS levels were quantified as the ratio of mean fluorescence intensity (MFI) of oxidized C11-BODIPY (FITC channel) to non-oxidized C11-BODIPY (PE channel) and normalized to corresponding DMSO-treated controls.

### GSH measurement

Total glutathione levels were measured using the GSH/GSSG-Glo™ Assay (Promega) according to the manufacturer’s instructions. Metastasis-derived Met (579DLN) and T_nonMet_ (579T1) cells were seeded at 1 × 10⁴ cells per well in 96-well plates and allowed to adhere overnight.

To assess baseline glutathione homeostasis, GSH/GSSG ratios were determined in untreated Met and T_nonMet_ cells under vehicle (DMSO) conditions. To evaluate the effect of artesunate on intracellular redox balance, cells were treated with artesunate (2.5 µM) for 24 h and GSH/GSSG ratios were measured.

To confirm glutathione depletion under conditions used for cell death analyses, total glutathione levels were assessed in Met and T_nonMet_ cell lines following treatment with artesunate, buthionine sulfoximine (BSO), or the combination for 6 h. In addition, to validate effective glutathione depletion prior to experimental metastasis assays, Met cells were treated with BSO (75 µM) for 24 h and total glutathione levels were measured.

Following treatment, culture medium was removed and replaced with 50 µL Total Glutathione Lysis Reagent. Plates were incubated on a shaker for 5 min, followed by addition of 50 µL Luciferin Generation Reagent and incubation for 30 min. Subsequently, 100 µL Luciferin Detection Reagent was added, and plates were incubated for 15 min.

Luminescence was measured using a BioTek Synergy H1 multimode plate reader. Glutathione levels were calculated according to the manufacturer’s protocol and normalized to DMSO-treated controls.

### Cell painting

Cell Painting was performed using an adapted protocol based on (Bray et al., 2016) and (Cimini et al., 2023). Cells were seeded in Millicell EZ 8-well glass slides (Millipore, cat. no. PEZGS0816) at a density of 5,000–10,000 cells per well in high-glucose DMEM and cultured at 37 °C, 5% CO₂. The next day cells were treated with artesunate 2.5 µM or with a vehicle for 24 h. For MitoTracker labeling, culture medium was replaced with a prewarmed medium containing MitoTracker Deep Red FM (Cell Signaling Technology, cat. no. 8778) at a final concentration of 500 nM, and cells were incubated for 30 min at 37 °C in the dark. Cells were then fixed by addition of 16% methanol-free paraformaldehyde (PFA) to a final concentration of 3.2% for 20 min at room temperature (RT). After one wash with 1× HBSS, cells were stained with WGA-Alexa Fluor 555 (Thermo Fisher Scientific, cat. no. W32464; 1.5 µg/mL) in 1% (w/v) BSA in HBSS for 7 min at RT, followed by two washes with HBSS. Cells were then permeabilized with 0.1% (v/v) Triton X-100 in HBSS for 15 min at RT, followed by two washes with HBSS.

Cells were stained for 30 min at RT in the dark with a cocktail containing Phalloidin-Alexa Fluor 568 (Thermo Fisher Scientific, cat. no. A12380; 5 U/mL), Concanavalin A-Alexa Fluor 488 (Thermo Fisher Scientific, cat. no. C11252; 100 µg/mL), and Hoechst 33342 (Thermo Fisher Scientific, cat. no. H3570; 5 µg/mL) in 1% (w/v) BSA in HBSS. Following three washes with HBSS, slides were mounted with ProLong Diamond Antifade Mountant (Invitrogen, cat. no. P36961) and stored in the dark until imaging. All experiments were performed in two independent biological replicates on different days.

Images were acquired on a ZEISS Axio Observer.Z1 widefield fluorescence microscope equipped with an ApoTome.2 module using a 63× objective across four channels: Hoechst (nucleus), Alexa Fluor 488 (endoplasmic reticulum), Alexa Fluor 555 (Golgi and plasma membrane), and Alexa Fluor 647 (mitochondria). Mitochondrial morphology was evaluated in five cells per cell line per condition per replicate, classifying mitochondria into different shapes as described by (Bulthuis et al., 2019). Two biologically independent experiments were performed.

### Anoikis assay

To assess anoikis resistance, Met (579DLN, 238N1, 482N1) and T_nonMet_ (579T1, 368T1, 394T4) cell lines were pretreated with artesunate (2.5 μM) for 72 h and subsequently seeded (2 × 10⁵ cells per well) into 6-well plates under either adherent (uncoated) or non-adherent conditions. Non-adherent conditions were generated by coating plates with poly (2-hydroxyethyl methacrylate) (PolyHEMA; 20 mg/mL in 95% ethanol; Sigma-Aldrich) to prevent cell attachment.

After 24 h, cells were harvested and stained with Annexin V for apoptosis analysis. Cell death was quantified by flow cytometry (BD FACSCelesta™, BD Biosciences) and analyzed using FlowJo® software. Anoikis sensitivity was determined by normalizing the percentage of Annexin V–positive cells under non-adherent conditions to adherent controls.

### Western blot

Met (579DLN, 238N1, 482N1) and T_nonMet_ (579T1, 368T1, 394T4) cell lines were seeded in 6-well plates at densities of 2 × 10⁵ cells per well for DMSO controls and up to 3 × 10⁵ cells per well for artesunate-treated samples to ensure comparable confluency at harvest. After 24 h, cells were treated with artesunate (2.5 μM) or DMSO for an additional 24 h.

Cells were washed twice with PBS and harvested by scraping. Cell pellets were washed and either processed immediately or stored at −20 °C until analysis. For protein extraction, pellets were lysed in an ice-cold lysis buffer and incubated on ice for 30 min. Lysates were clarified by centrifugation at 4 °C, and supernatants were collected. Protein concentrations were determined using the BCA Protein Assay Kit (Thermo Fisher Scientific) according to the manufacturer’s instructions.

Equal amounts of protein were mixed with 4× Laemmli buffer containing 10% β-mercaptoethanol, heated at 95 °C for 5 min, and separated by SDS-PAGE using 4–15% Mini-PROTEAN TGX precast gels (Bio-Rad). Proteins were transferred onto PVDF membranes using a semi-dry transfer system. Membranes were blocked in 5% BSA in TBS-T and incubated overnight at 4 °C with primary antibodies against GPX4 (Abcam, ab125066, RRID: AB_10973901) and GAPDH (Cell Signaling Technology, 14C10, RRID: AB_561053).

After washing, membranes were incubated with HRP-conjugated anti-rabbit secondary antibody (Cell Signaling Technology; 7074S, RRID: AB_2099233, 1:10,000) for 1 h at room temperature. Protein bands were visualized using Clarity™ Western ECL substrate (Bio-Rad) and detected with a ChemiDoc imaging system (Bio-Rad). Experiments were performed in three independent replicates.

### In vivo

All animal experiments were carried out following ethical guidelines and approved by the Landesamt für Verbraucherschutz und Ernährung (LAVE). Mice were housed in individually ventilated cages under specific pathogen-free conditions with *ad libitum* access to food and water. Animals were maintained on a 12 h light/12 h dark cycle with controlled temperature and humidity. All animal experiments and data analyses were conducted without blinding, and no prior power analysis was performed to determine sample size.

### *In vivo* multiplexed screening for metastatic seeding

Metabolic drug multiplexed screening was performed as described by (Grüner et al., 2016). Briefly, 579DLN (Met) and 579T1 (T_nonMet_) cell lines were stably barcoded and treated with a metabolic compound library for 72 h *in vitro*. Following treatment, cells from each plate were trypsinized, pooled, and prepared as a suspension of approximately 1 × 10⁶ cells in PBS, which was injected via the tail vein into syngeneic 129/B6 F1 mice. Mice of 8 weeks or older and both sexes were used. The remaining pooled cells from each plate (pre-injection pools) were centrifuged, and pellets were stored at −80 °C until DNA isolation. In parallel, cell confluency was measured using the NyOne system on three additional plates with identical drug layouts and concentrations to ensure that observed effects were not attributable to reduced *in vitro* cell growth.

Forty-eight hours post-injection, lungs were harvested and processed according to the digestion protocol. The resulting cell pellet was resuspended in 800 μL of sorting buffer, and barcoded tumor cells were enriched by flow cytometry based on Tomato fluorescence. Sorted cells were centrifuged at 300 × g for 5 min, and pellets, together with pre-injection pools, were used for DNA extraction, PCR amplification, and next-generation sequencing (NGS). Sequencing data were analyzed as described by (Grüner et al., 2016).

### Pre-treatment rescue experiment

For the *in vivo* rescue experiment, female immunocompetent 129/B6 F1 mice aged ≥9 weeks were used (control, artesunate [ART], and ART+NAC: *n* = 6; NAC: *n* = 5 per group). The Met (579DLN) cell line was pretreated *in vitro* for 24 h with artesunate (2.5 μM), N-acetylcysteine (NAC; 10 mM), vehicle (1% DMSO), or the combination of artesunate and NAC. Following treatment, cells were trypsinized, centrifuged, and resuspended at 1 × 10⁷ cells/mL in PBS, and 100 μL (1 × 10⁶ cells) was injected intravenously via the tail vein into each mouse.

Forty-eight hours post-injection, lungs were harvested and processed according to the digestion protocol. Tumor burden was quantified by flow cytometric analysis of Tomato⁺ cells, expressed as the percentage of Tomato⁺ cells in lung tissue.

### Intravenous injection of BSO-pretreated cells

Immunocompetent 129/B6 F1 mice (both male and female, aged ≥ 13 weeks) were used (control *n* = 6; BSO *n* = 7 per group). The Met (579DLN) cell line was pretreated *in vitro* with BSO (75 μM) or vehicle control for 24 h. Following treatment, cells were harvested and 1 × 10⁶ cells were injected intravenously via the tail vein into each mouse. Forty-eight hours post-injection, lungs were harvested and processed according to the digestion protocol. Tumor burden was quantified by flow cytometric analysis of Tomato⁺ cells and expressed as the percentage of Tomato⁺ cells in lung tissue.

### *In vivo* subcutaneous injections with artesunate treatment

In this study, NOD SCID gamma (NSG) mice (both male and female, aged ≥ 8 weeks) were used and randomly assigned to experimental groups (control *n* = 11; artesunate-treated *n* = 12). For the spontaneous metastasis model, 5 × 10⁴ Met (579DLN) cells were resuspended in 100 μL PBS and kept on ice until injection. Immediately prior to injection, 100 μL Matrigel® (Corning, 356231) was added, and 200 μL of the 1:1 cell suspension/Matrigel mixture was injected subcutaneously into each flank. Tumor growth and animal health were monitored regularly. Nine days post-injection, when tumors became palpable, mice were randomized into treatment groups. Control mice received daily intraperitoneal (i.p.) injections of saline supplemented with 2.5% sodium bicarbonate (NaHCO₃; Sigma-Aldrich, S6014), while the treatment group received artesunate (Hölzel-Biotech, MOLN-M16406) at 100 mg/kg. After 16 days of treatment, mice were sacrificed in accordance with approved animal welfare protocols.

Tumors, lung lobes, and selected organs (liver and kidneys) were imaged from both sides using a Stereo Discovery V8 microscope (Zeiss). Tumors and selected lung lobes were preserved for histological analysis. For metastasis quantification, lung lobes were processed using a digestion protocol. Lung tissue was minced and digested for 30 min at 37 °C in digestion medium containing 10% trypsin-EDTA (0.25%; Thermo Fisher Scientific, 25200056), 10% collagenase IV (Gibco, 17104019), and dispase (Corning, 354235) in HBSS without Ca²⁺ and Mg²⁺ (Gibco, 14175095). The reaction was quenched with Leibovitz’s L15 medium supplemented with 10% FBS and DNase I (final concentration 3.75 μL/mL from a 5 mg/mL stock; Sigma-Aldrich, 11284932001), followed by filtration through a 40 μm cell strainer.

Cells were centrifuged, and red blood cells were lysed using ammonium–chloride–potassium (ACK) buffer. The remaining cell pellet was resuspended in PBS containing 5 μg/mL DAPI (Thermo Fisher Scientific, D21490) and analyzed by flow cytometry (BD FACSCelesta™, BD Biosciences) for Tomato⁺ cells.

Experiments were performed in two independent biological replicates, and data were normalized to the respective control group within each experiment prior to analysis.

### Imaging and histology

#### Histology and immunohistochemistry

Lung and tumor tissue samples were fixed in 4% paraformaldehyde (PFA) overnight and embedded in paraffin. Sections were cut and stained with hematoxylin and eosin (H&E) using standard methods. For immunohistochemical analysis, antigen retrieval was performed in 1 mM citrate buffer (pH 6.0), followed by incubation with an anti-Ki67 primary antibody (Abcam, ab15580, RRID: AB_443209). Whole-slide images were acquired using an Axio Scan.Z1 slide scanner (Zeiss), and quantification was performed using Orbit Image Analysis (https://www.orbit.bio). A supervised classifier was trained to distinguish Ki67-positive nuclei (DAB, brown), Ki67-negative nuclei (hematoxylin, blue), and background tissue. For each mouse, two tumor sections were analyzed, with ten ROIs of identical size randomly selected per tumor (twenty ROIs per mouse). The Ki67 index was calculated as the proportion of Ki67-positive nuclei relative to total nuclei, and mean values were determined per section for statistical analysis.

### Transcriptomics and bioinformatics

#### Bulk RNA sequencing

Bulk RNA sequencing was performed on T_nonMet_ (579T1) and Met (579DLN) cells following treatment with vehicle (0.5% DMSO) or artesunate (2.5 μM) for 48 h.

Three biological replicates of T_nonMet_ (579T1) and Met (579DLN) cells were treated with artesunate (2.5 µM) or DMSO (0.5% DMSO) for 48 h and frozen. Frozen cell pellets were sent to Singleron Technologies GmbH (Cologne, Germany), where RNA isolation and bulk transcriptome sequencing were performed.

In brief, total RNA was isolated using the RNeasy Mini Kit (Qiagen, cat. No. 74106). Subsequently, RNA quantity and integrity were assessed using Qubit^TM^ RNA High Sensitivity (HS) Assay Kits (Invitrogen^TM^, cat. no. Q10211) and High Sensitivity RNA ScreenTape (Agilent, cat. no. 5067-5579). Purified RNA was processed according to the instructions of the AccuraCode^®^ RNA-Seq kit (Singleron Biotechnologies GmbH, cat. no. 10710174) with the following modifications. Briefly, polyadenylated mRNA from different samples was captured using paramagnetic AccuraCode bulk beads (Singleron Biotechnologies GmbH, mat. no. 1200030080). During reverse transcription, the mRNA was converted into cDNA and labeled with sample-specific barcodes and unique molecular identifiers (UMI). The labeled cDNA was pooled, amplified and used to construct RNA-seq libraries according to manufacturer’s instructions (Singleron Biotechnologies GmbH, AccuraCode^®^ RNA-Seq kit, cat. no. 10710174). The libraries were sequenced on an Illumina NovaSeq X instrument using a paired-end 150-bp approach. Raw sequencing reads were processed into gene expression matrices using CeleSCOPE v2.0.7 (RRID:SCR_023553) multi_bulk_rna with customized chemistry (https://github.com/singleron-RD/CeleScope; Singleron Biotechnologies GmbH). Sequencing reads were aligned to the Mus musculus reference genome (Ensembl release 99) using STAR (RRID:SCR_004463), and gene-level counts were generated with featureCounts (RRID:SCR_012919). KEGG (RRID:SCR_012773) pathway enrichment analysis was performed using the clusterProfiler package (Yu et al., 2012). Differentially expressed genes with an adjusted *P* value (FDR) < 0.05 and an absolute log2 fold change > 1 were subjected to over-representation analysis against the mouse KEGG database (*mmu*) using Benjamini– Hochberg multiple testing correction. Gene symbols were converted to Entrez gene identifiers using the org.Mm.eg.db annotation database prior to enrichment analysis.

Baseline differential gene expression analysis between untreated 579DLN (Met) and 579T1 (T_nonMet_) cells was performed using DESeq2 (RRID:SCR_000154). Normalized gene expression values were used for visualization of glutathione-related gene expression.

Volcano plots were generated from DESeq2 differential expression results by plotting log2 fold change against the −log10 adjusted *P* value. Genes with an adjusted *P* value (FDR) < 0.05 and an absolute log2 fold change > 1 were considered significantly differentially expressed and colored according to the direction of regulation. Selected genes of interest were manually annotated.

### Microarray Data Analysis of redox gene expression

Microarray gene expression data were obtained from the publicly available Gene Expression Omnibus (GEO) (RRID:SCR_005012) dataset GSE26874 (Winslow et al., 2011), comprising murine lung adenocarcinoma cell lines derived from primary tumors or metastases generated by conditional activation of *Kras*^G12D^ and deletion of *Trp53*. Processed and normalized expression data generated using the Affymetrix Mouse Genome 430 2.0 Array platform were downloaded directly from GEO, and no additional normalization was applied. Expression levels of selected redox-related genes (*Slc7a11*, *Gss*, *Gpx1*, *Gclc*, *Gclm*, *Gsr*, *Slc25a39, Atf4, Idh1*) were compared between primary tumor–derived and metastasis-derived cell lines. Median expression values were calculated per group, and effect size was defined as the difference in median expression (Δ = Metastasis − Primary). Genes were ranked according to Δ values for visualization.

### Single-cell RNA sequencing

Publicly available single-cell RNA sequencing data from lung adenocarcinoma patients (GEO accession GSE131907 (Kim et al., 2020)) were used for analysis. Malignant epithelial cells were selected according to the published cell type annotations provided with the dataset. Gene expression values for glutathione-related genes were extracted from malignant epithelial cells originating from primary lung tumors, brain metastases, and lymph node metastases. For patient-level analyses, the mean expression of each gene was calculated across all malignant epithelial cells from each individual patient. Primary tumors were compared with metastatic lesions comprising brain and lymph node metastases. Differences in patient-level mean gene expression were assessed using a two-sided Wilcoxon rank-sum test. False discovery rates (FDRs) were calculated using the Benjamini–Hochberg method. Data processing, statistical analyses, and visualization were performed in R (version 4.5.3).

### Statistics

Statistical analyses were performed as specified in the figure legends, and *P* values are indicated in the figures. Graphs were generated using GraphPad Prism (version 7.05) (RRID:SCR_002798), R (version 4.4.1 and 4.5.3), and Python (RRID:SCR_024202) within the Jupyter Notebook (version 7.0.8) in an Anaconda environment.

## RESOURCE AVAILABILITY

### Lead contact

Further information and requests for resources and reagents should be directed to the lead contact, Barbara M. Grüner.

## KEY RESOURCES TABLE

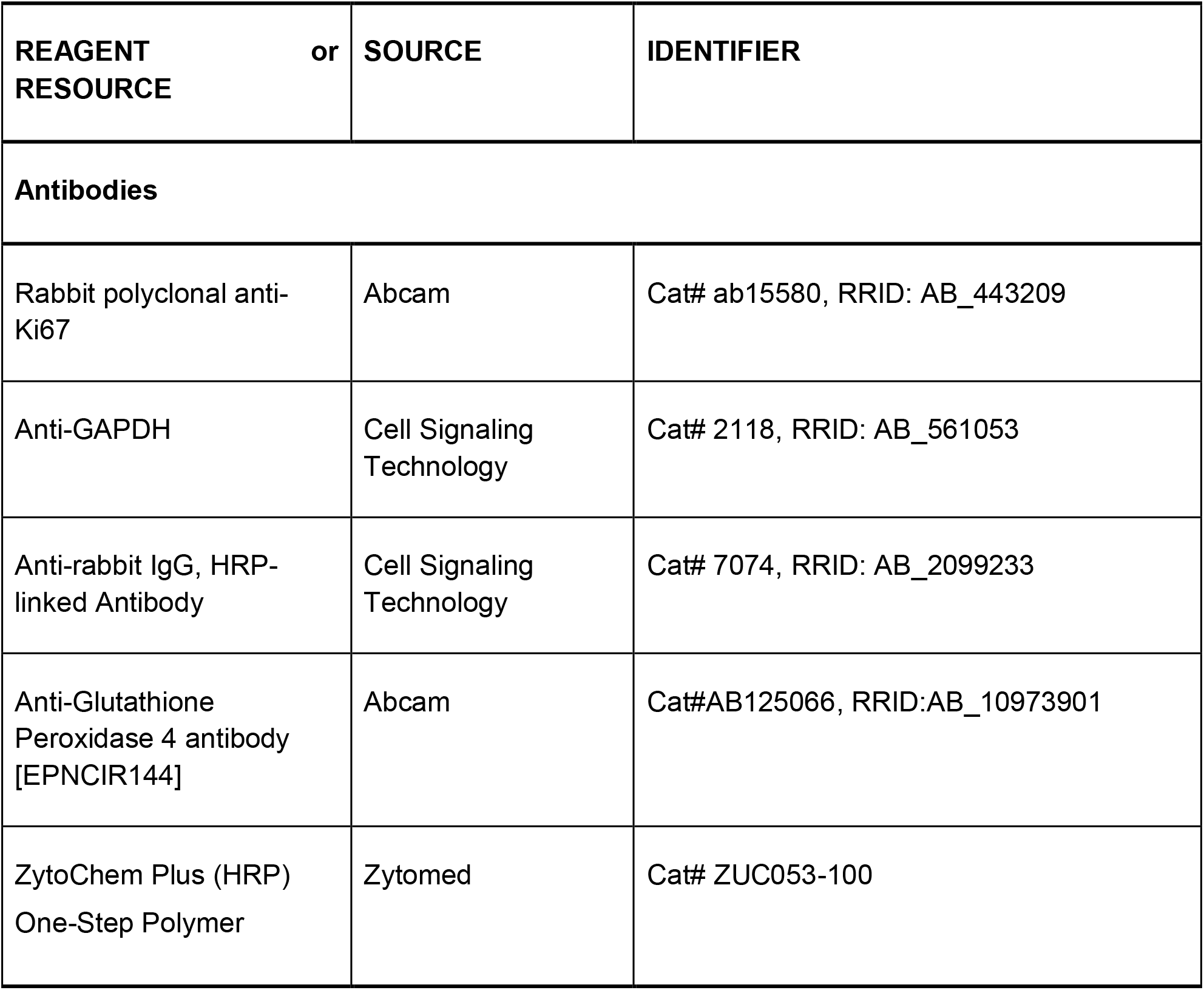
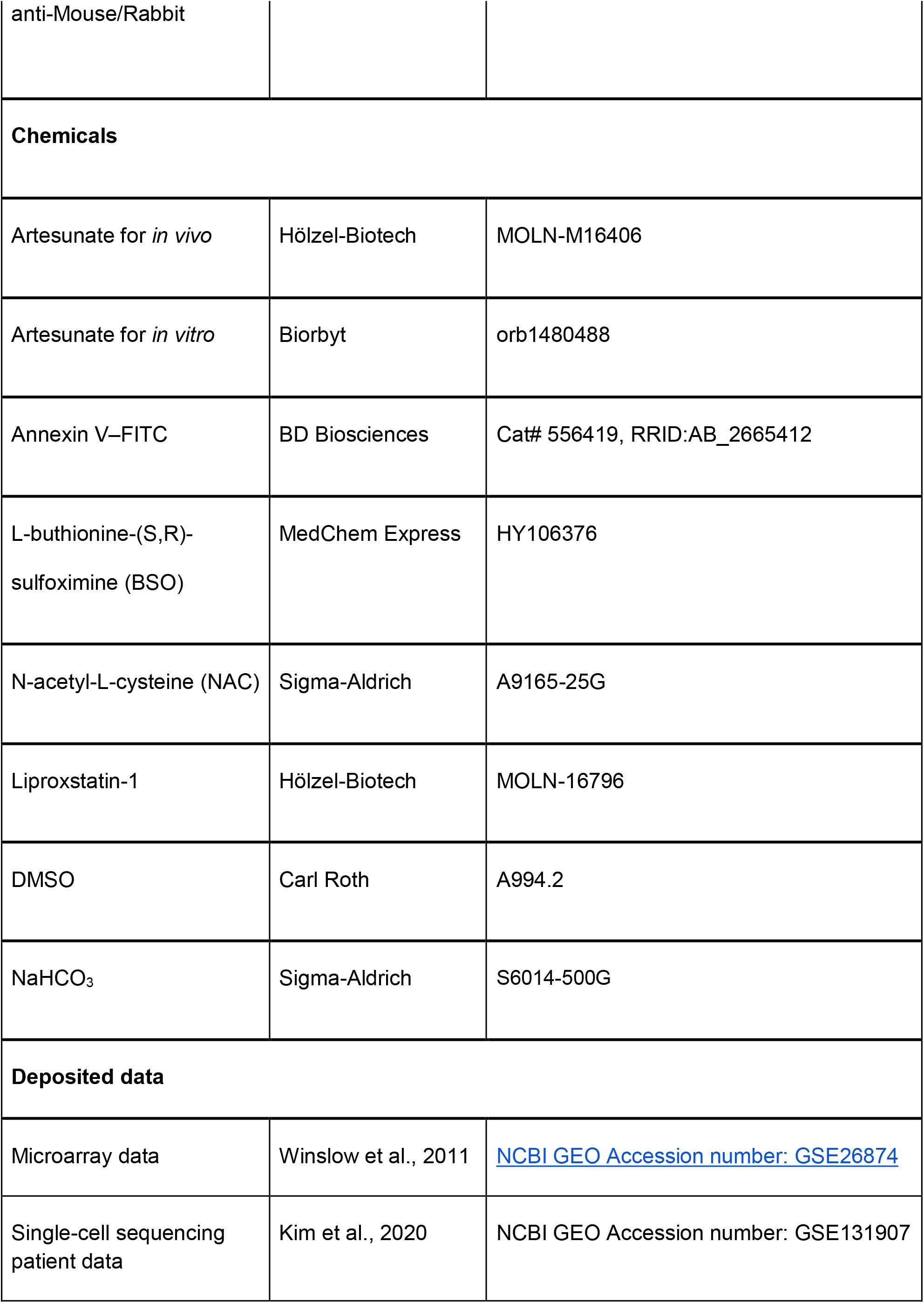
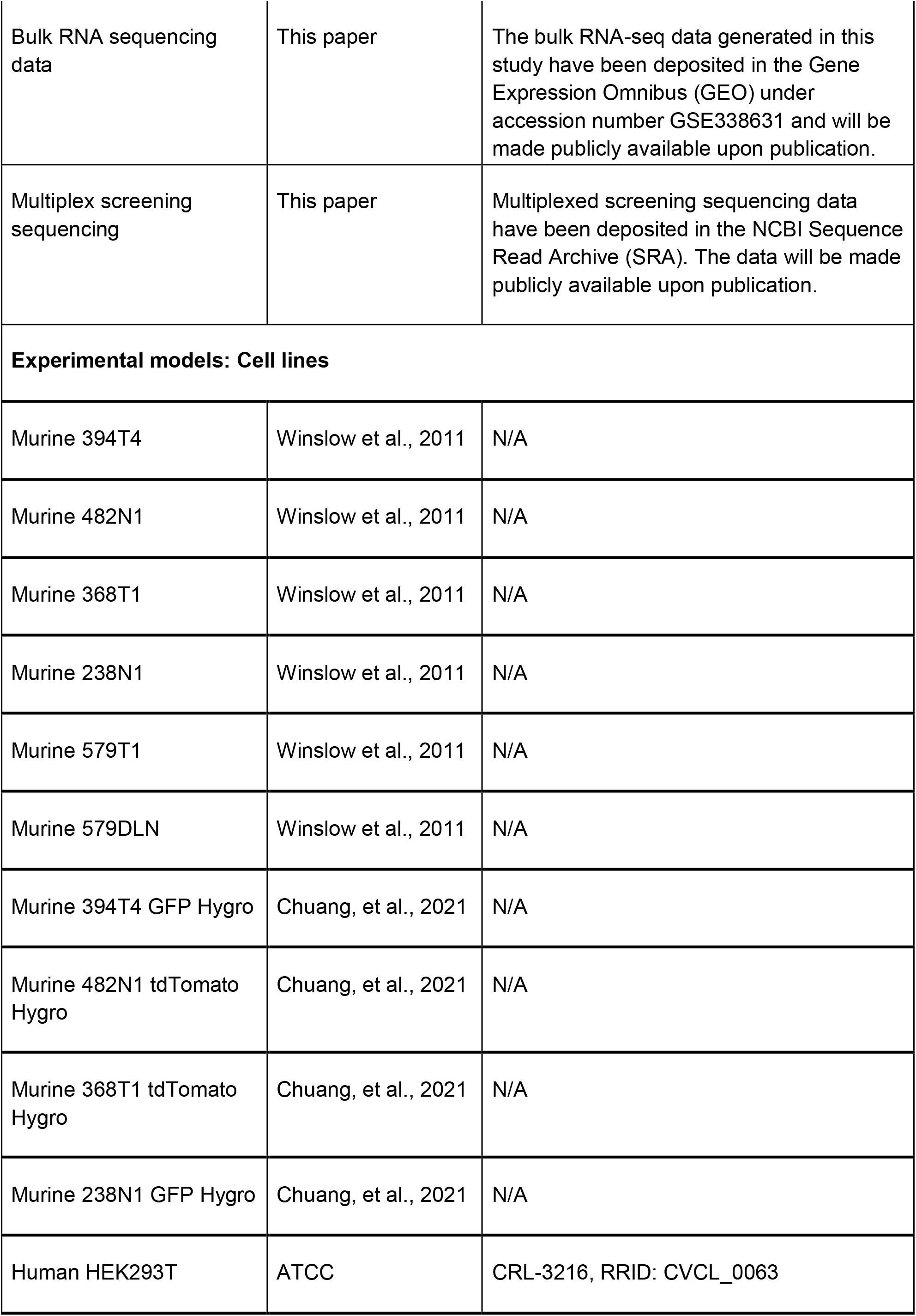
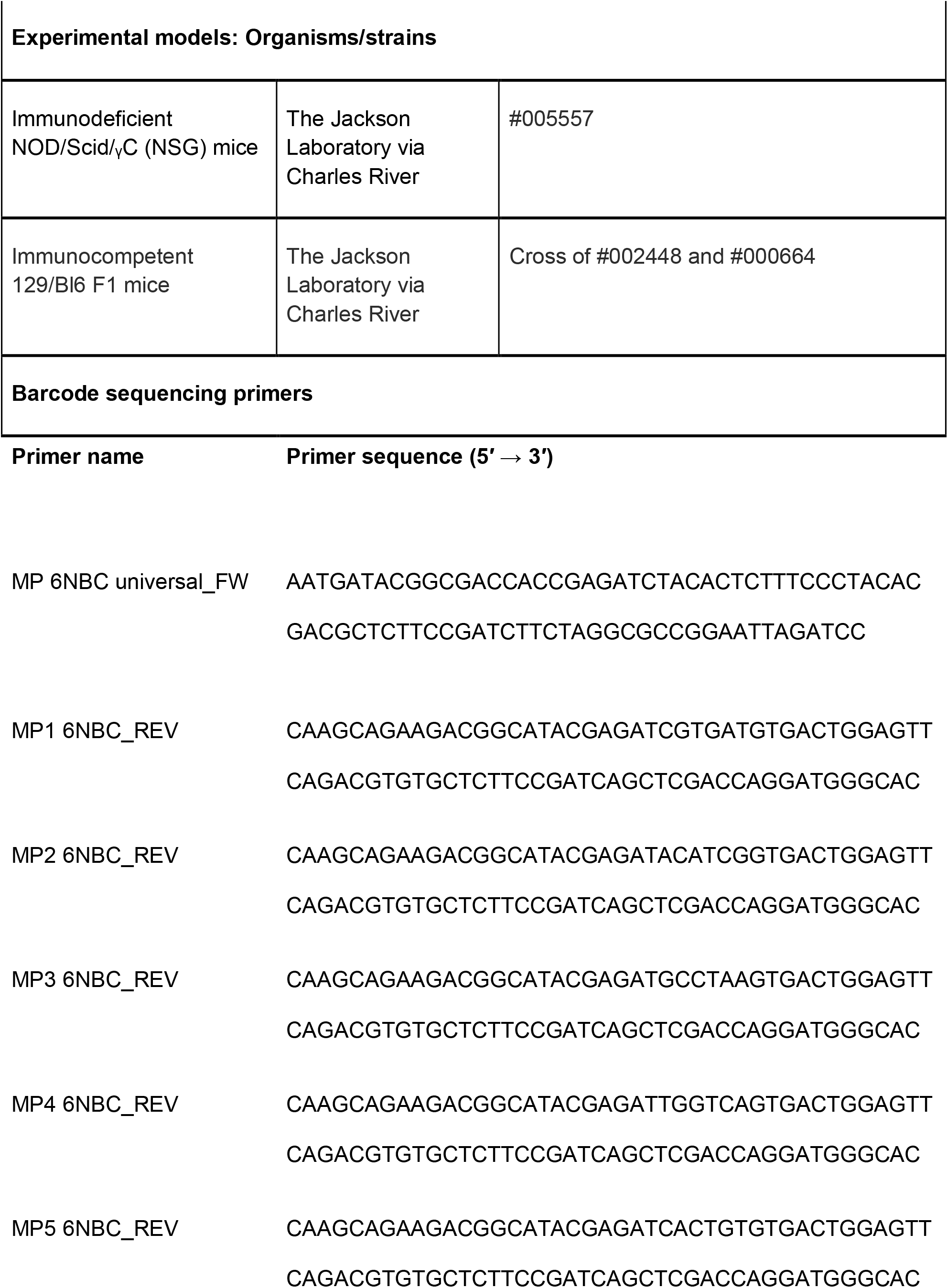
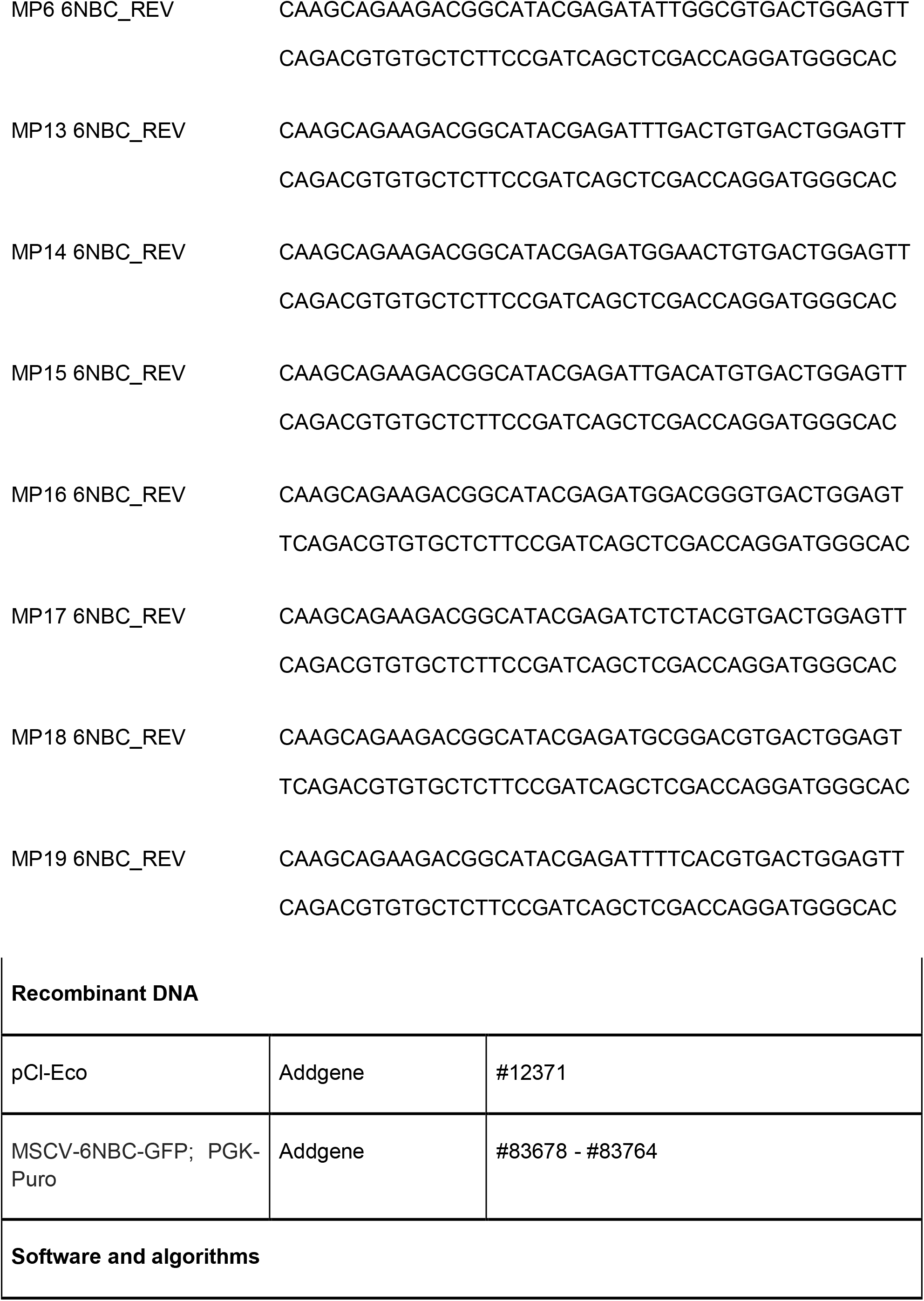
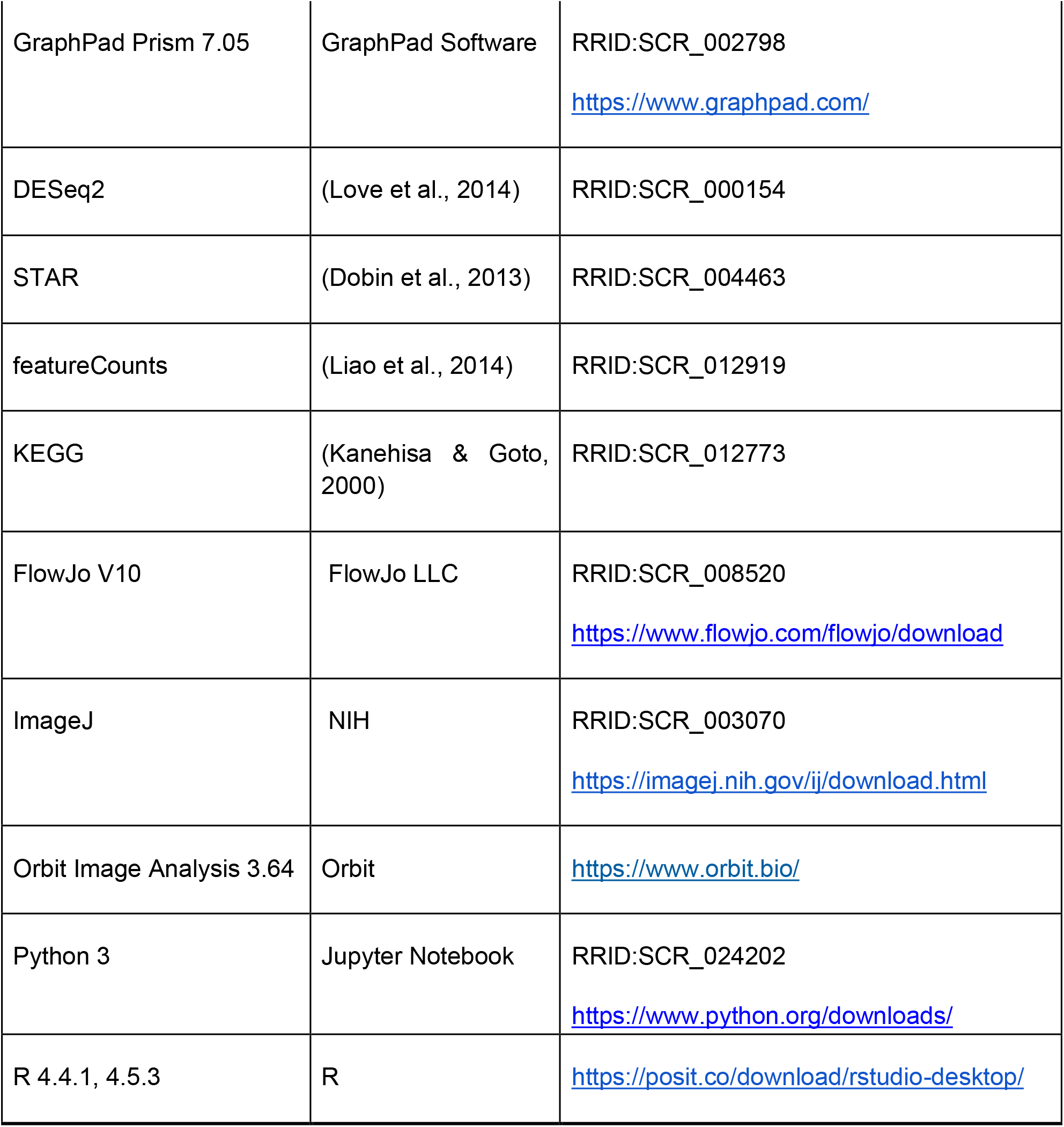

## ACKNOWLEDGMENTS

We thank Eva Hahn and Nancy Meyer for excellent technical assistance, and the members of the Grüner, Schramm, Grünwald, and Siveke labs for helpful comments and discussions. We also thank the central animal facility of the University Hospital Essen (ZTL) for their support. This work was supported by the German Cancer Aid (DKH - 70114056) and the German Research Foundation (DFG CRU337 - 405344257) to B.M.G., as well as the BMFTR (DKTK-ED03) to V.S. and J.C.B.

## AUTHOR CONTRIBUTIONS

Conceptualization: P.M., B.M.G.

Methodology: P.M., V.S., J.C.B.

Formal analysis: P.M.

Investigation: P.M., S.N., P.F., V.S., J.C.B., A.B.

Validation: P.M., B.M.G.

Resources and concepts: J.C.B., J.P.F.A., A.S.

Writing – original draft: P.M., B.M.G.

Writing – review & editing: all authors Funding acquisition: B.M.G.

