## Supplementary Figures for "Artesunate Exploits Glutathione-Dependent Redox Vulnerability in Metastatic Lung Adenocarcinoma"

Munteanu *et al.* 2026\_Supplementary Figure 1

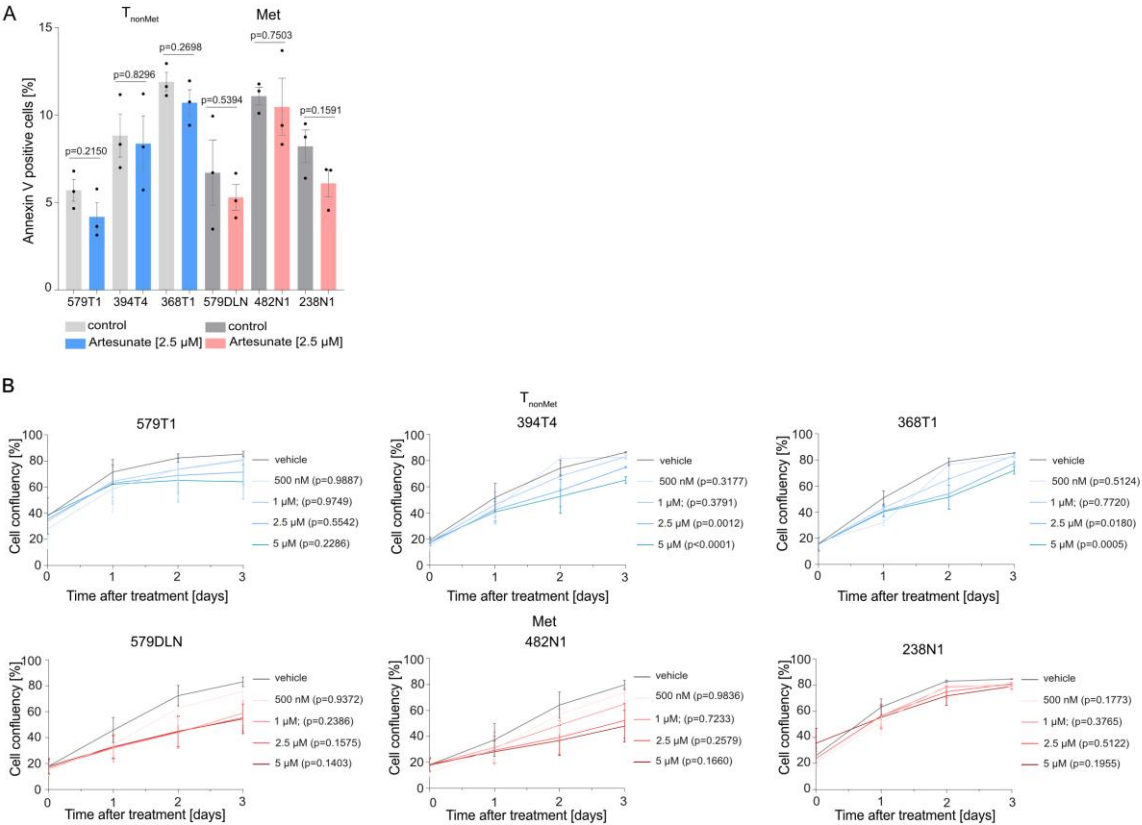

### Supplementary Figure 1

**(A)** Annexin V–positive apoptotic cells following treatment with artesunate (2.5 μM). Quantification shows the percentage of Annexin V–positive cells across three independent experiments ($n = 3$). Data are presented as mean ± SEM. Statistical significance was assessed using a two-tailed unpaired Welch's $t$-test.

**(B)** Cell growth of lung cancer cell lines was assessed by longitudinal confluency measurements following treatment with increasing concentrations of artesunate (0.5, 1, 2.5, and 5 μM) using the NyOne live-cell imaging system. Data are shown as mean ± SEM from $n = 3$ independent experiments. Statistical significance was assessed using Dunnett's multiple comparisons test.

Munteanu *et al.* 2026\_Supplementary Figure 2

A

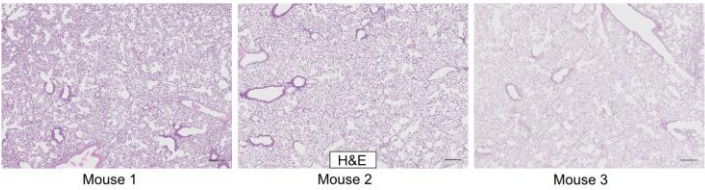

B

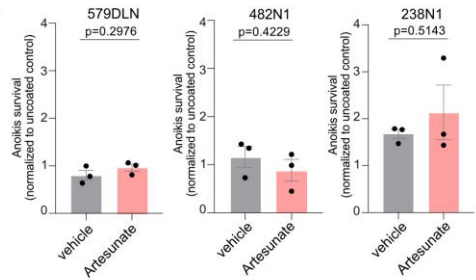

### Supplementary Figure 2

**(A)** Representative H&E images of lungs of the mice at treatment initiation ($n = 3$).

Scale bars represent 200 $\mu\text{m}$.

**(B)** Metastatic cell lines (238N1, 482N1, and 579DLN) were treated with artesunate for 72 h, followed by analysis of Annexin V–positive cells seeded in poly-HEMA-coated wells. Data were normalized to the corresponding uncoated (poly-HEMA-free) condition. Data represents SEM from three independent experiments ($n = 3$). Statistical significance was assessed using a two-tailed unpaired Welch's $t$-test.

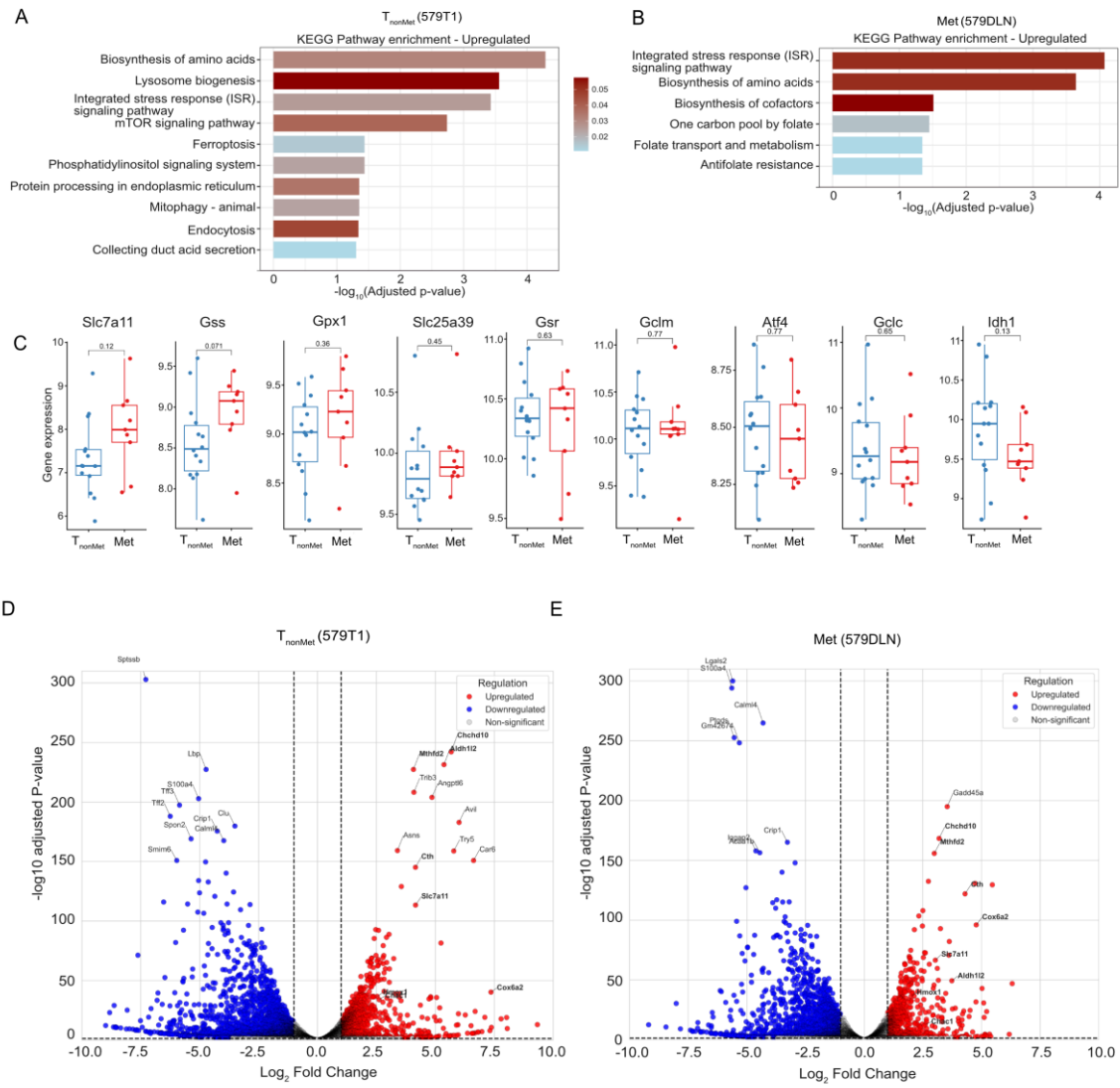

#### Supplementary Figure 3

**(A-B)** KEGG pathway enrichment analysis in 579T1 and 579DLN cells following 48 h treatment with 2.5 μM artesunate. RNA sequencing was performed using three independent biological replicates per condition. Bubble plots showing upregulated pathways based on bulk RNA sequencing. The x-axis represents the $-\log_{10}(\text{adjusted p-value})$ of pathway enrichment, with larger values indicating greater statistical significance. Bar color indicates the gene ratio, representing the proportion of differentially expressed genes overlapping with each pathway.

**(C)** Expression of glutathione- and redox-related genes in murine primary lung adenocarcinomas ($n = 14$) and metastatic lesions ($n = 9$) from the GEO dataset GSE26874 (Winslow et al., 2011). Differences in gene expression between primary tumors and metastatic lesions were analyzed using two-sided unpaired Welch's $t$-test.

**(D-E)** Volcano plots showing transcriptomic changes in 579T1 and 579DLN cells after 48 h treatment with 2.5 μM artesunate. Differential gene expression was assessed using DESeq2. Genes with $|\log_2 \text{fold change}| \geq 1$ and adjusted p-value ($P_{\text{adj}}$) $\leq 0.05$ were considered significantly differentially expressed. Upregulated genes are shown in red, downregulated genes in blue, and non-significant genes in grey. Volcano plots are shown for 579T1 (D) and 579DLN (E) cells.

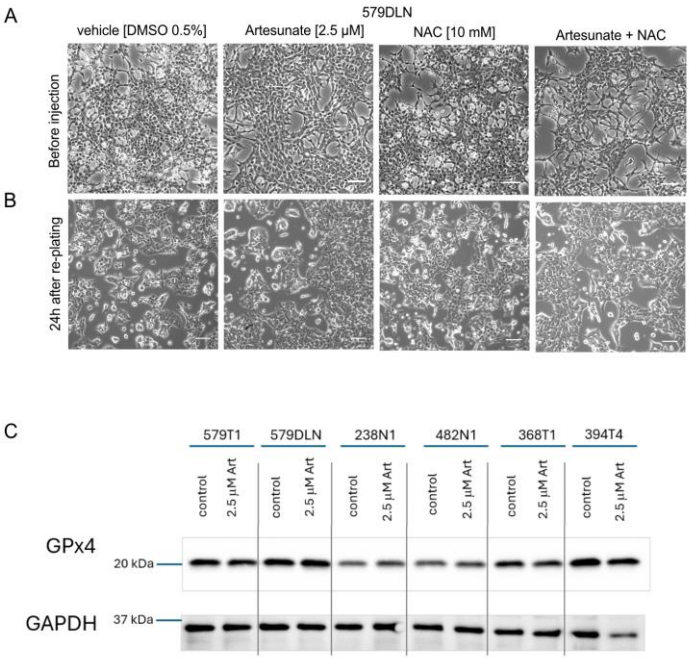

#### **Supplementary Figure 4**

**(A)** Morphology of cells before and after intravenous (IV) injection under indicated treatment conditions. Cells were treated with vehicle (DMSO 0.5%), artesunate (2.5 μM), N-acetylcysteine (NAC, 10 mM), or the combination of artesunate and NAC for 24 h prior to intravenous injection. A fraction of the same cell suspension used for injection was returned to culture, re-plated, and imaged at 24 h **(B)** post-plating to assess cell recovery and morphology following injection-related handling. Representative images of cells immediately before injection (upper panel) and 24 h after replating (lower panel). Scale bars, 50 μm.

**(C)** Western blot analysis of GPX4 protein levels following treatment with 2.5 μM artesunate for 24 h. GAPDH served as a loading control. One representative blot from three independent experiments is shown.

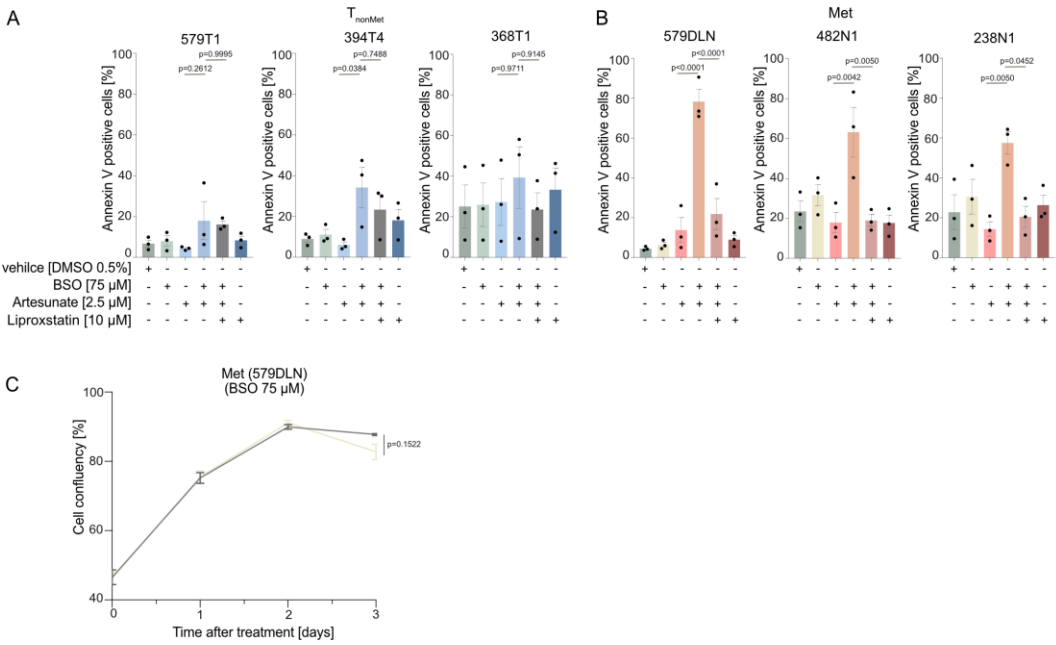

#### Supplementary Figure 5

**(A-B)** Flow cytometric quantification of Annexin V–positive cells in T<sub>nonMet</sub> (A) and Met (B) cell lines 72 h after treatment with DMSO, artesunate (2.5 μM), artesunate + BSO (75 μM), or artesunate + BSO + liproxstatin-1 (10 μM). Early and late apoptotic populations were combined as total Annexin V–positive cells. Data represent mean ± SEM from three independent experiments. Statistical significance was assessed using a one-way ANOVA followed by Tukey's multiple comparisons test.

**(C)** Cell growth of 579DLN cells following treatment with BSO (75 μM). Cell confluency was monitored using the NyOne live-cell imaging system over 72 h and is presented as percentage confluency over time. Error bars represent SEM of technical replicates. Statistical significance at the 72 h endpoint was assessed using a two-tailed unpaired Welch's *t*-test.
